# Bayesian inference of transcriptional branching identifies regulators of early germ cell development in humans

**DOI:** 10.1101/167684

**Authors:** Christopher A. Penfold, Anastasiya Sybirna, John Reid, Aracely Castillo Venzor, Elena Drousioti, Yun Huang, Murray Grant, Lorenz Wernisch, Zoubin Ghahramani, M. Azim Surani

**Affiliations:** Wellcome/CRUK Gurdon Institute, University of Cambridge; Wellcome/MRC Cambridge Stem Cell Institute, University of Cambridge; Physylogy, Development and Neuroscience Department, University of Cambridge; MRC Biostatistics Unit, University of Cambridge; Department of Engineering, University of Cambridge; School of Life Sciences, University of Warwick

**Author notes:** These authors should be considered joint first authors (Christopher A. Penfold, Anastasiya Sybirna).

## Abstract

During embryonic development, cells undertake a series of fate decisions to form a complete organism comprised of various cell types, epitomising a branching process. A striking example of branching occurs in humans around the time of implantation, when primordial germ cells (PGCs), precursors of sperm and eggs, and somatic lineages are specified. Due to inaccessibility of human embryos at this stage of development, understanding the mechanisms of PGC specification remains difficult. The integrative modelling of single cell transcriptomics data from embryos and appropriate *in vitro* models should prove to be a useful resource for investigating this system, provided that the cells can be suitably ordered over a developmental axis. Unfortunately, most methods for inferring cell ordering were not designed with structured (time series) data in mind. Although some probabilistic approaches address these limitations by incorporating prior information about the developmental stage (capture time) of the cell, they do not allow the ordering of cells over processes with more than one terminal cell fate. To investigate the mechanisms of PGC specification, we develop a probabilistic pseudotime approach, branch-recombinant Gaussian process latent variable models (B-RGPLVMs), that use an explicit model of transcriptional branching in individual marker genes, allowing the ordering of cells over developmental trajectories with arbitrary numbers of branches. We use first demonstrate the advantage of our approach over existing pseudotime algorithms and subsequently use it to investigate early human development, as primordial germ cells (PGCs) and somatic cells diverge. We identify known master regulators of human PGCs, and predict roles for a variety of signalling pathways, transcription factors, and epigenetic modifiers. By concentrating on the earliest branched signalling events, we identified an antagonistic role for FGF receptor (FGFR) signalling pathway in the acquisition of competence for human PGC fate, and identify putative roles for PRC1 and PRC2 in PGC specification. We experimentally validate our predictions using pharmacological blocking of FGFR or its downstream effectors (MEK, PI3K and JAK), and demonstrate enhanced competency for PGC fate *in vitro*, whilst small molecule inhibition of the enzymatic component of PRC1/PRC2 reveals reduced capacity of cells to form PGCs *in vitro*. Thus, B-RGPLVMs represent a powerful and flexible data-driven approach for dissecting the temporal dynamics of cell fate decisions, providing unique insights into the mechanisms of early embryogenesis. Scripts relating to this analysis are available from: https://github.com/cap76/PGCPseudotime

## 1 Introduction

During embryogenesis, individual cells undertake a series of cell fate decision to form a complete embryo comprised of myriad cell types. Each cell fate decision can be thought of in terms of a bifurcation (Poincaré 1885), with the expression levels of key marker genes diverging between the two cell fates, epitomising a branching process. Reciprocal behavior is encountered in recombination processes, where two or more statistical processes converge, such as when two or more intermediate cell types share a common terminal fate. A key challenge is to infer the mechanisms and inductive signals of these decision-making processes by identifying the ordering of bifurcations of individual genes using single-cell RNA-sequencing.

A striking example of transcriptional branching occurs in early human development, when the inner cell mass (ICM) of the blastocyst segregates into hypoblast and epiblast, with the latter subsequently differentiating into ectoderm, mesoderm, and endoderm during the process of gastrulation (Irie, Tang, and Azim Surani 2014). At around this time, circa weeks 2-3 in humans, primordial germ cells (PGC), the embryonic precursors of gametes, are also specified (Irie et al. 2015; Kobayashi et al. 2017). Later, at weeks 5-6, specified PGCs undergo comprehensive epigenetic reprogramming that includes almost complete erasure of DNA methylation marks throughout the genome, save for a few escapee regions (Guo et al. 2015; Tang et al. 2015; Gkountela et al. 2015). During this period, PGCs also proliferate and migrate towards the genital ridges where, after colonising the gonads, they begin sexually dimorphic programs of gametogenesis. As precursors of the germline, PGCs are ultimately responsible for passing on all genetic and epigenetic information to the next generation, and any aberrant development has the potential to lead to infertility or cancers of the germ line.

Despite recent advances in understanding of PGC development (Irie et al. 2015; Tang et al. 2015), the mechanisms of human PGC development remain poorly understood. Current understanding, based on studies in mouse and pig embryos, suggests that two signalling pathways cooperate to allow PGC fate: WNT signalling renders epiblast cells competent to respond to BMP2/4 resulting in the specification of a founder PGC population in the posterior epiblast (Ohinata et al. 2009; Aramaki et al. 2013; Kobayashi et al. 2017). While mouse PGC development has been reasonably well characterised, the specification and development of human PGCs remains only partially understood, primarily due to inaccessibility of early human embryos. Crucially, recent studies have shown that *SOX17* and *PRDM1* are key regulators of PGC fate in humans (Irie et al. 2015; Tang et al. 2015), making their specification distinct from that in mice, which involves the combined action of *Prdm1*, *Prdm14*, and *Tfap2c* (Magnusdottir et al. 2013; Nakaki et al. 2013). Due to these differences, along with notable divergence in embryo morphology (Irie, Tang, and Azim Surani 2014), *in vitro* derivation of human PGC-like cells (PGCLCs) from human pluripotent stem cells has emerged as a model to examine the earliest mechanisms regulating hPGC development (Sasaki et al. 2015; Irie et al. 2015). Interestingly, human ESCs in conventional cultures have low competence to form PGCLCs, but hESCs grown in a specially formulated “competent” medium gain competence for PGC fate and respond to BMP signalling, giving rise to PGCLCs that presumably resemble pre-migratory *in vivo* PGCs (Irie et al. 2015).

The integrative analysis of single cell RNA-seq data from pre-implantation embryos, specified PGCs, and appropriate *in vitro* models, should provide unique opportunities to dissect the dynamics of early PGC cell fate decisions. Such analysis requires that cells be correctly ordered along a continuous developmental trajectory. However, most approaches for pseudotemporal ordering of scRNA-seq datasets rely on manifold learning: that is, a preliminary dimensionality reduction, with cells ordered over the reduced dimensional space, typically by utilising curve fitting or graph-theoretic approaches (Trapnell et al. 2014; Bendall et al. 2014; Marco et al. 2014; Ji and Ji 2016; Setty et al. 2016). Such approaches do not generally account for uncertainty in the ordering, nor do they usually provide an interpretable relationship between the inferred pseudotime and chronological time. The latter limitation is compounded when inferring pseudotimes for datasets with multiple branches: if one branch has fewer observations, or else a period of quiescence, the branch will often be artificially truncated compared to the others. Such truncation can be undesirable when investigating complex developmental programs, where signalling from adjacent tissues influence cell fate decisions, making it necessary to place branches on consistent timeframes. Finally, in the past, scRNA-seq datasets tended to be of low temporal resolution, albeit with observations in many cells, which reflect a continuum of developmental states (Yan et al. 2013; Guo et al. 2015; Petropoulos et al. 2016; Borensztein et al. 2017; Huang et al. 2017). However, due to decreased costs, scRNA-seq datasets are now routinely generated over finely resolved time series, including the early stages of embryogenesis (Yan et al. 2013; Guo et al. 2015; Petropoulos et al. 2016; Borensztein et al. 2017; Huang et al. 2017; Ibarra-Soria et al. 2018; Han et al. 2018) and during PGC development (Guo et al. 2015; Li et al. 2017). While individual populations of cells associated with the different stages still reflect a continuum of states, with some degree of overlap between stages, the existence of a well-defined capture time provides highly informative prior information about the ordering of cells, which most approaches are incapable of utilising.

To address these various limitations, and investigate the mechanisms of PGC specification, we develop a probabilistic approach to pseudotemporal ordering. Our approach incorporates prior information about developmental stage of cells (capture time), and uses an explicit model of branching at the level of marker genes, providing an interpretable model of cell fate decision making designed specifically for time-series scRNA-seq data. Using our model, we combine data from preimplantation embryos, PGCs, somatic tissues, and human embryonic stem cells (hESCs), to dissected the transcriptional program and signalling pathways of human PGC competence, specification, and development. Our analysis highlights the importance of known PGC genes, and suggested several novel regulators. Analysis suggested the importance of polycomb repressive complexes 1 and 2 (PRC1 and PRC2) in PGC specification, and small-molecule inhibition of PRC1/2 enzymatic activity was shown to reduce PGC specification using human *in vitro* models of PGC specification. Crucially, identification of the earliest branching pathways highlighted putative roles for FGF receptor (FGFR) signalling in the acquisition of competence for human PGC fate, which was experimentally validated *in vitro*. Indeed, pharmacological blocking of FGFR or its downstream effectors (MEK, PI3K and JAK) enhanced the competency for PGC fate *in vitro*. Thus, our approach can inform genetic and signalling perturbations for cell fate decisions.

## 2 Results

Early human PGC development remains underexplored, since the use of human embryos around the time of specification is limited by ethical and practical considerations. This has necessitated the development of *in vitro* models of PGC development to bridge the gap in understanding at key developmental stages (Irie, Sybirna, and Surani 2018). Statistical approaches are required to correctly leverage multiple *in vivo* and *in vitro* datasets to identify priority targets for more focused experiments.

To separate out the developmental trajectories of PGCs from those of gonadal somatic cells, we ordered single cells along a two-component branching process, with informative priors placed over the pseudotimes centered on the cells’ developmental stage (Reid and Wernisch 2016) (Figure 1; Supplementary Materials Section 1). We used scRNA-seq data from hPGCs and age-matched neighbouring somatic tissues from weeks 4 through to 19 (Guo et al. (2015); GEO GSE63818), as well as pre-implantation embryos (oocytes, zygotes, two-cell, four-cell, morula and blastocyst-stage cells), and conventional hESCs at passages 0 and 10 (Yan et al. (2013); GEO GSE36552). A summary of the various cell types is included in Supplementary Table 1.

**Figure 1:**
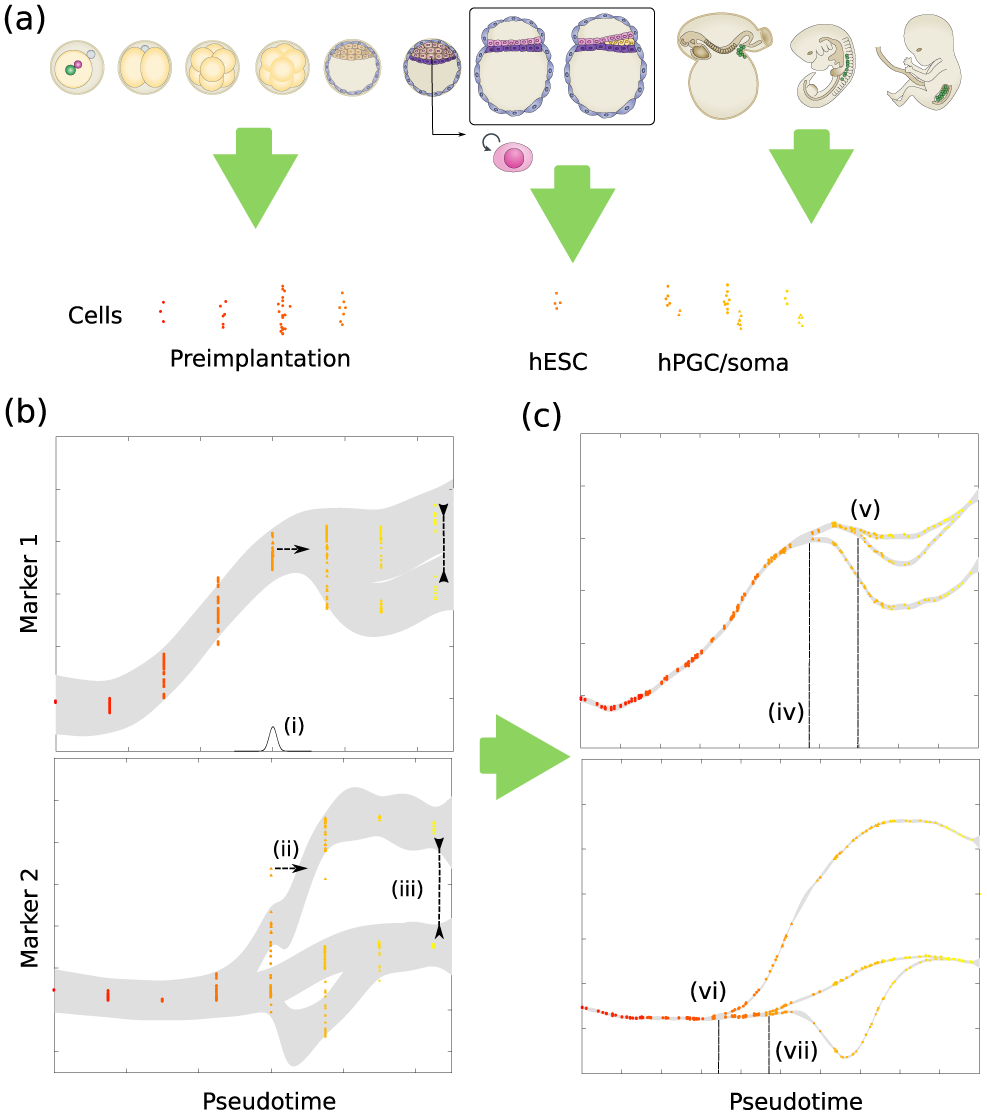
(a) Here we performed an integrative analysis of human cells from preimplantation embryos, hESCs, hPGCs and age matched soma. Due to the structured nature of the data, we decided to take a Bayesian approach to the analysis allowing us to take advantage of useful prior information, such as capture time. (b) Initially data is ordered by capture time over a range of marker genes. Using an iterative (Monte Carlo) approach, we permute cells along the pseudotime by perturbing a subset of cells along the x-axis (ii) or by allowing cells to swap branch assignments (iii). Following each perturbation, the marginal likelihood or “evidence” can be computed, and used to determine whether to accept or reject the proposed move. (c) After many iterations, cells are ordered along a branching process that reflects the developmental progression of the system In this case, we can identify a sequential branching of marker genes 1 and 2, with the first branch (iv, vi), and a subsequent branching and recombination (v, vii). By comparing the pseudotime of these branching events between genes (compare iv with vi, and v with vii) we can identify the earliest events in cell fate decisions and the developmental hierarchy.

Based on preliminary benchmarking experiments, ordering of cells was based on the expression levels of 44 marker genes identified from Irie et al. (2015), which included PGC, pluripotency, mesoderm, and endoderm markers. To evaluate the accuracy of our approach we used unlabelled data i.e., assuming the branch-labels and capture-times for hESCs and blastocyst cells were unknown variables to be assigned during inference. Our approach correctly placed the unlabelled late-blastocyst cells between morula-stage and week 4 (Wk4) cells (Figure 2(a)). Furthermore, whilst hESCs are derived from the ICM of blastocysts, our approach placed them between day 6 blastocysts and Wk4 cells (PGCs and soma), albeit with some degree of overlap. This suggests that hESCs cultured under these conditions are developmentally more advanced than cells of the ICM, consistent with conventional hESCs sharing characteristics with the post-implantation epiblast (Tang et al. (2016); Figure 2(a), inset).

**Figure 2:**
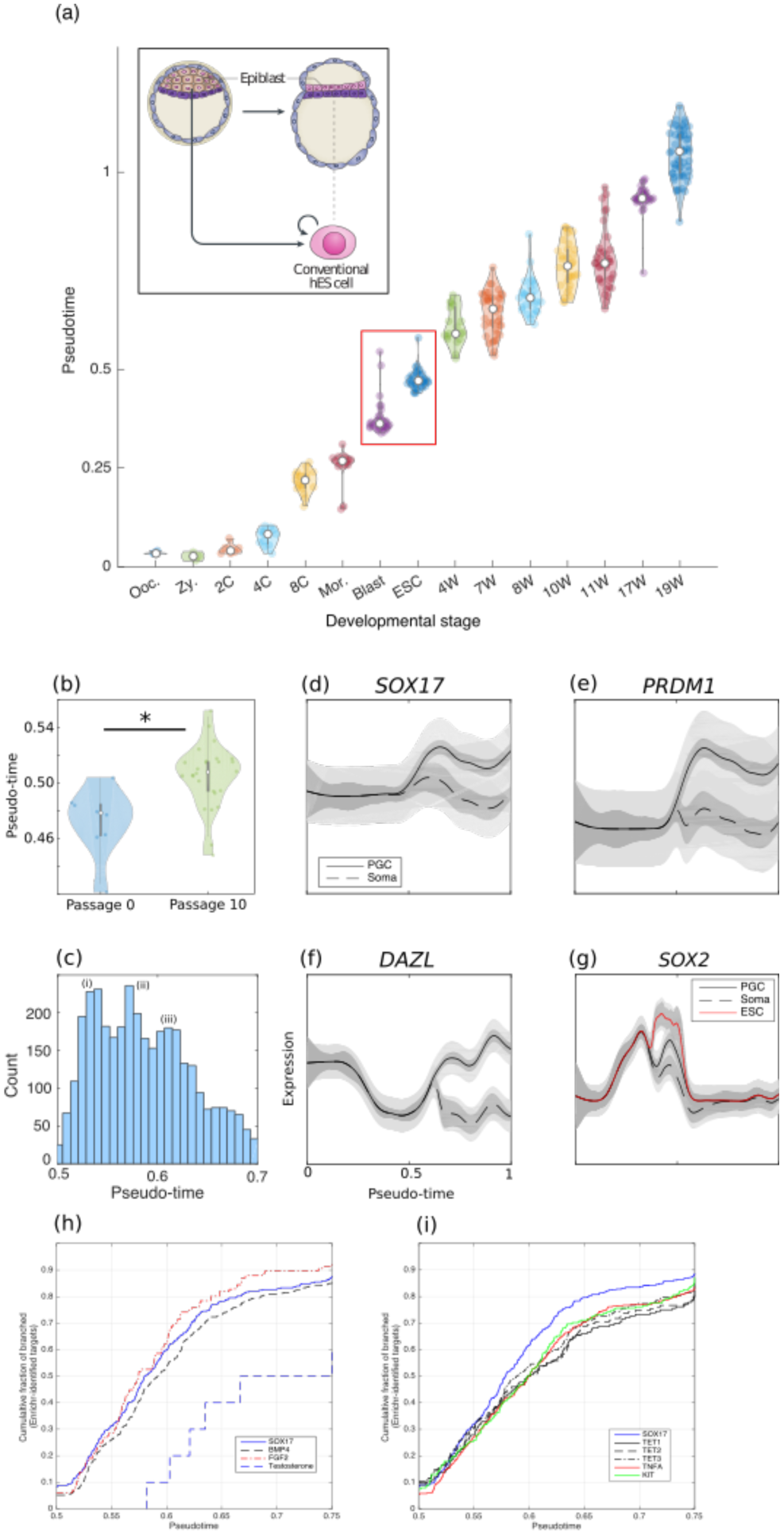
**(a)** Inferred pseudotemporal ordering of individual cells over a two-component branching component correctly identifies the developmental ordering of blastocyst stage cells and ESCs, suggesting hESCs are developmentally more advanced than blastocyst stage. **(b)** ESCs at passage 10 appear to be developmentally more advanced than at earlier passages, having statistically later pseudotimes. **(c)** Histogram of branching time (soma versus PGCs) indicates a multimodal response. **(d, e, f)** Key PGC regulators SOX17 and PRDM1 branch early in the pseudotime series, prior to late PGC markers such as DAZL. **(g)** SOX2, a pluripotency and neuronal cell fate gene, branches between ESCs and the inferred in vivo dynamics. **(h)** Identified targets of SOX17 and BMP4 from perturbation studies in ESCs show early branching between PGCs and soma. **(i)** TNF response appears to be concomitant with genes associated with epigenetic reprogramming and the migratory phase of PGC development.

To further evaluate the performance of our approach, we calculated: (i) the Pearson’ correlation coefficient between the inferred pseudotime, and the developmental stage; and (ii) a branch discrepancy metric between somatic and PGC lineages:

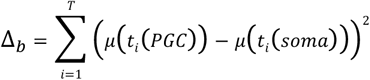

Where *μ*(*t_i_*(*PGC*)) is the mean inferred pseudotime of PGC cells at developmental stage *i*. For comparison, we ordered cells using established methods including Monocle2 (Qiu et al. 2017), TSCAN (Ji and Ji 2016), Wishbone (Setty et al. 2016), SCUBA (Marco et al. 2014), SLICER (Welch, Hartemink, and Prins 2016) and GrandPrix (Ahmed, Rattray, and Boukouvalas 2018). Results are summarised in Supplementary Table 2/Supplementary Figure 3.

**Figure 3:**
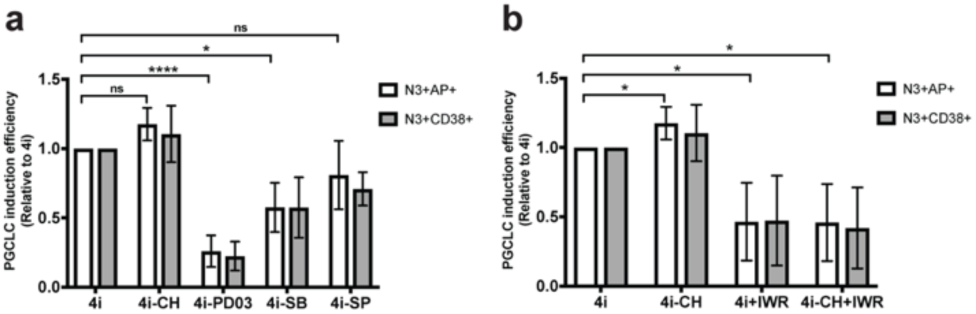
Quantification of PGCLC induction efficiency from hESCs grown in indicated conditions relative to 4i hESCs. PGCLCs induction efficiency was defined as the percentage of live NANOS3-tdTomato/AP (N3+AP+) or NANOS3-tdTomato/CD38-double positive cells. Data are shown as mean ± SD of 2 or 4 independent experiments. * p ≤ 0.05, **** P ≤ 0.0001, ns: not significant (p > 0.05), Holm-Sidak t-test (on relative frequency of N3+AP+ cells). **(a)** MEK and p38 inhibitors withdrawal from 4i medium decreases PGCLC competence of hESCs. **(b)** Inhibition of WNT signalling by small molecule inhibitor IWR reduces PGCLC competence.

Overall, our approach offered the best performance, with Pearson’ correlation of 0.96 for the PGC branch and 0.95 for the soma branch, with low branch-discrepancy metric (∆_*b*_= 0.004) indicating good alignment between the two lineages. Monocle2, TSCAN, and GrandPrix all performed well at ordering cells albeit with slightly lower correlation coefficient, with GrandPrix also doing a good job of aligning the two branches (∆_*b*_= 0.02). Whilst other approaches showed a general ability to separate out pre-implantation cells from PGC or soma, they did not necessarily place cells along a continuous trajectory, and did not appear able to align different branches.

To evaluate the effect of more complex branching structures, we also ordered cells along a 3-component branching process, explicitly modelling where ESCs and soma diverged from the PGC trajectory. Here we additionally investigated the effect of using increased numbers of genes in the algorithm: using 44 marker genes; 87 marker genes; and the top 101 most varied genes as an unbiased alternative to our marker-based strategy. In all cases the B-RGPLVM ordered cells along a continuous developmental trajectory, with high correlation between pseudotime and capture time, and low branch-discrepancy metric. Performance did not appear to increase as the number of observed genes was increased, suggesting that cells could be accurately ordered with around 40 marker genes, in agreement with preliminary analysis using other datasets.

### 3.2.2. Inferring branching structure on a genome scale

The preliminary ordering of cells suggested that key PGC makers SOX17 and PRDM1 branched earlier than late markers such as DAZL. Although the data was not resolved enough to distinguish the order of branching between SOX17 and PRDM1, the posterior distribution suggested that SOX17 branched prior to late PGC markers such as DAZL with >90% certainty.

To identify novel regulation on a genome scale, we subsequently used independent B-RGP regression (Penfold et al. 2018) to infer branching on a gene-by-gene basis, conditional on the estimated pseudotime. Whilst this allowed inference of branching for all expressed genes, this increased scalability comes at a loss of information about posterior distribution of pseudotimes, and does not quantify the uncertainty in ordering. For each gene, we explicitly assumed one of three models: (i) somatic cells branched from the base process (the hPGCs trajectory), with hESCs and hPGCs following an identical process (see Figure 2(d, e, f)); (ii) both soma cells and hESCs branched from the main process, with hESCs later recombining towards Wk4 PGCs (Figure 2(g)); and (iii) all processes were identically distributed (no branching). Similarly to our approach in (Penfold et al. 2018) we used the Bayesian information criterion (BIC) to determine the branching structure for each gene (Supplementary Section 3). The number of genes assigned to each group is indicated in Supplementary Figure 4(a), and shows that most genes were non-branching i.e., not differentially expressed.

**Figure 4.**
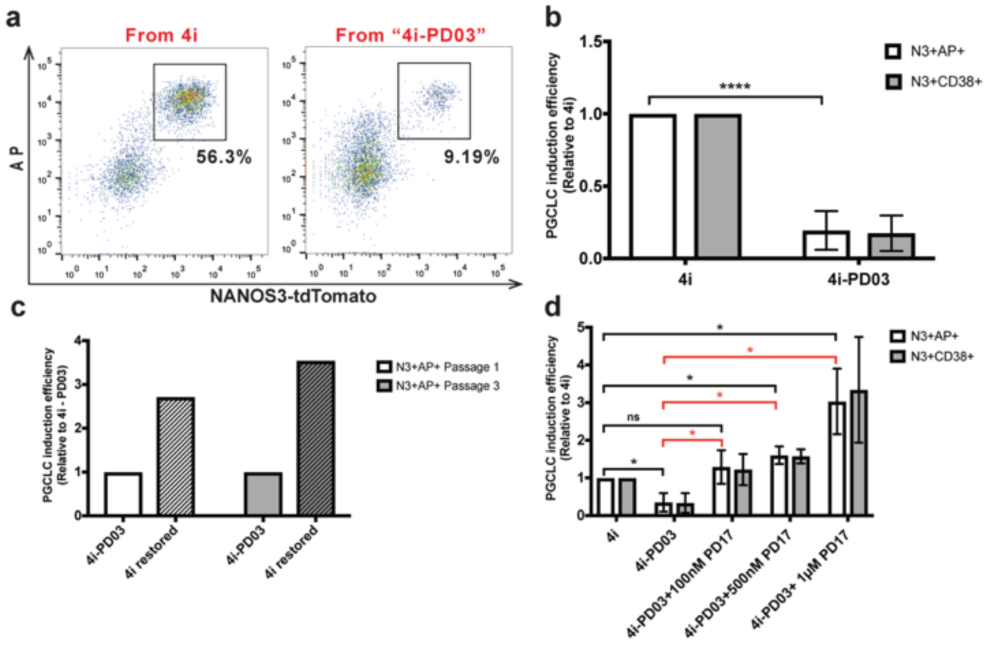
FGFR-MEK signalling is a negative regulator of human PGCLC competence. **(a)** MEK inhibitor withdrawal from 4i medium decreases PGCLC competence of hESCs Representative flow cytometry plots of EBs derived from 4i and “4i-PD03” hESCs. **(b)** Quantification of PGCLC induction efficiency from “4i-PD03” relative to 4i hESCs. Data are shown as mean ± SD of 9 independent experiments. **** p ≤ 0.0001, Holm-Sidak t-test (on relative frequency of N3+AP+ cells). **(c)** “4i-PD03” differentiation defect can be rescued by reintroducing PD03. hESCs grown in “4i-PD03” for >10 passages were transferred to complete 4i medium (with PD03) for 1 or 3 passages and subjected to PGCLC induction. Induction efficiency is shown relative to “4i-PD03”-derived PGCLCs; n=1. **(d)** hESCs grown with FGFR inhibitor PD17 show enhanced PGCLC competence. Data are shown as mean ± SD of 2 or 3 independent experiments. * p ≤ 0.05, ns: not significant (p > 0.05), Holm-Sidak t-test (on relative frequency of N3+AP+ cells). Red lines and asterisks refer to comparison of “4i-PD03” to other conditions.

#### 3.1 In vivo dynamics of human PGC development

In total, 3,930 genes were identified as being up-regulated in hPGCs versus soma, with 1,355 down-regulated. It is likely that these genes are involved in a broad range of biological processes, including the acquisition of competence for PGC fate, PGC specification and maintenance, as well as epigenetic reprogramming, migration, and gametogenesis. We therefore performed a preliminary GO and KEGG pathway analysis using a permissive p-value (p<0.1, Bonferroni corrected hypergeometric test), to identify biological processes that branched early in development (Supplementary File 1). Our analysis revealed several early terms related to BMP and WNT signalling, with subsequent terms associated with epigenetic reprogramming, proliferation and cell migration, and later terms relating to testosterone signalling, meiosis, and gametogenesis which, together, reflect the expected progression of hPGC development (Lawson et al. 1999; Ohinata et al. 2009; De Felici 2013; Leitch, Tang, and Surani 2013; Kobayashi et al. 2017).

A histogram of the time of branching between PGCs and soma (Figure 2(c)) shows a multi-modal distribution, with most responses occurring after blastocyst-stage, but prior to week 4, consistent with the expected timing for PGC specification at around weeks 2-3 of development (see e.g., (De Felici 2013; Tang et al. 2016)). Importantly, even when using a point estimate for pseudotime, the two key regulators of human PGC fate, *SOX17* and *PRDM1* (Irie et al. 2015), were still shown to branch early, prior to late PGC markers such as *DAZL* (Figure 2(d, e, f)). In addition to regulating human, but not mouse, PGC fate, *SOX17* is a known endoderm TF in mice and humans (Kanai-Azuma et al. 2002; Seguin et al. 2008). It is therefore possible that a subset of its target genes are shared between human endoderm and PGC lineages. To test this, we looked at genes that were differentially expressed in hESCs following overexpression of *SOX17* compared to parental hESC lines (Seguin et al. (2008); GSE10809). Genes that branched between PGCs and soma were statistically enriched for genes that were differentially expressed in response to *SOX17* overexpression (p<1×10^-35^, hypergeometric test using Enrichr). Reanalysis of this microarray data showed that common genes included the PGC regulator *PRDM1*, known to act downstream of *SOX17* (Irie et al. 2015). Crucially, the overlapping targets of *SOX17* were, themselves, shown to branch early compared to the dataset as a whole (p<1×10^-7^, Kolmogorov–Smirnov test), and earlier than testosterone signalling, involved in later male germ cell development (Figure 2(h)).

To reveal additional regulatory mechanisms that could be contributing to PGC specification and maintenance, we used Enrichr (Chen et al. 2013; Kuleshov et al. 2016) to find transcriptional signatures present within our set of branched genes, and used a Kolmogorov-Smirnov (KS) test to identify those signatures that branched significantly earlier than average branching times in the dataset. That is, we looked for early and statistically significant overlaps between genes that branched (PGCs versus soma) and those that were up-regulated or down-regulated in the literature in a variety of species and cell types following knockout or overexpression studies, upon treatment with hormones, growth factors, and cytokines, or were overrepresented in Encode ChIP datasets or the ChEA database. Although none of the Enrichr perturbations were performed directly in PGCs, and results might therefore be contextually very different to our system, the presence of an early and statistically robust transcriptional signature is worthy of further investigation. Furthermore, we envisage that in this way it may be possible to identify signalling molecules secreted from surrounding tissues and therefore not branching in the examined transcriptome itself.

In agreement with the early branching of genes with BMP-related GO/KEGG terms around the time of PGC specification, comparison with the Enrichr database showed a strong overlap of branched genes with those differentially expressed in hESCs following treatment with BMP4 (p<5×10^-14^, Enrichr SG). The identified targets of BMP4 branched early in the pseudotime series compared to branch times overall (p<5×10^-4^, KS-test; Figure 2(h)), reflecting the known role of BMP4 as a PGC-inductive signal (Lawson et al. 1999). We further identified the transcriptional signatures of several putative BMP and WNT effector genes, including Parathyroid Hormone Like Hormone (*PTHLH*, adjusted p<5e^-5^, Enrichr SG), which increases mesenchymal cells’ responsiveness to BMP4 (Hens et al. 2009), and has been implicated in the emergence of germ cell cancers (Sandberg, Meloni, and Suijkerbuijk 1996; Mostert et al. 1998). Other signatures included X-linked inhibitor of apoptosis (*XIAP*, p<5×10^-26^, Enrichr SG) which, besides modulating BMP and WNT signalling, has also been associated with seminomas, cancers originating from PGCs (Kempkensteffen et al. 2007; Oosterhuis and Looijenga 2005). Within branched genes, we also noted strong signatures of genes involved in canonical WNT signalling, including both *CTNNB1* (p<5×10^-18^, Enrichr SG; p<0.001, KS-test), *GSK3β* (p<5×10^-6^, Enrichr Kinase; p<0.05, KS-test), as well APC Membrane Recruitment Protein 1 (*AMER1,* p<1e-11, Enrichr SG; p<5×10^-6^, KS-test), and its target, Wilms Tumor 1 (*WT1*, p<5×10^-7^, Enrichr SG). These findings are in keeping with WNT and BMP serving as major PGC induction signals and thus prove the efficacy of our framework.

Amongst the early enriched gene signatures was overexpression of the transcription factor *NANOG* (p<0.05 KS-test). In mouse *in vitro* models NANOG is sufficient to directly induce PGC-like cell fate by binding and activating enhancers of *Prdm1* and *Prdm14* (Murakami et al. 2016), although loss of function studies have yielded equivocal results as to its requirement for mouse PGC fate (Chambers et al. 2007; Yamaguchi et al. 2009; Carter et al. 2014). NANOG is highly expressed in human PGCs and PGCLCs, but, unlike in mouse cells, its overexpression does not induce PGCLCs (Kobayashi et al. 2017). Nevertheless, we identified a number of putative NANOG binding sites within the 10kb flanking regions of *PRDM1* using FIMO (Supplementary File 2), suggesting a possible functional role. It would be of interest to test the involvement of NANOG in hPGCLC fate using an inducible knockout.

In addition to picking up known PGC regulators, we were also able to identify other pathways potentially involved in human PGC fate, including a putative role for tumour necrosis factor (TNF) signalling in PGC development, based on the enrichment of KEGG term ‘TNF signalling pathway’, and the TNF transcriptional signature (p<1×10^-10^, Enrichr SG). Targets of TNF were shown to branch later than BMP4/SOX17-responsive genes, and around the same time as genes that branched in response to perturbations of *TET1*, *TET2*, and *TET3* (Figure 2(i)). This places the timing of TNF-signalling roughly in concordance with epigenetic reprogramming in proliferating, migratory PGCs. Although TNFA has roles in apoptosis, previous studies in mouse models suggest it can stimulate proliferation of pre-gonadal PGCs in culture (Kawase et al. 1994; Makoolati, Movahedin, and Forouzandeh-Moghadam 2016). Whilst *TNF,* itself, was not identified amongst the branched genes, and its expression appears to be generally low in PGCs (Tang et al. 2015), one of its receptors, *TNFRSF1A*, was found to be branched, and we cannot exclude that TNF is secreted from surrounding tissues. Indeed, *TNFA* is expressed in Schwann cells (Wagner and Myers 1996), and TNF signalling might therefore support PGCs proliferation and survival as they migrate along the nerve fibres to the genital ridges (Mollgard et al. 2010).

#### 3.1.1 Branched genes were enriched for targets of polycomb repressive complexes 1 and 2 (PRC1 and PRC2)

Branched genes were also enriched for the signature of epigenetic and chromatin modifiers, including the histone demethylase *KDM3A* (p<1×10^-5^, Enrichr SG; p<0.001, KS-test), methyltransferase *DNMT1* (p<5×10^-5^, Enrichr; NS, KS-test), and dioxygenases *TET1 (*p<1×10^-4^, Enrichr SG)*, TET2* (p<5×10^-4^, Enrichr SG), and *TET3* (p<5×10^-5^, Enrichr SG), which contribute to PGC DNA demethylation via conversion of the modified genomic base 5-methylcytosine (5mC) into 5-hydroxymethylcytosine (5hmC) (Ito et al. 2010). In general, genes associated with perturbations of epigenetic and chromatin modifiers appeared to branch later than genes associated with perturbations of WNT and BMP signalling, in line with existing experimental evidence (Tang et al. 2015). Amongst the related enriched GO terms, we noted several related to cAMP activity, and the top 10 enriched motifs between genes up-regulated in PGCs and the ChEA database included several cAMP modulators, including CREB1 (p<5×10^-44^, Enrichr ENCODE) and CREM (p<5×10^-90^, Enrichr ChEA), consistent with studies in mice that identify roles for cAMP signalling in mouse PGC proliferation and epigenetic reprogramming (Ohta et al. 2017).

The polycomb group (PcG) of proteins are a diverse and evolutionary conserved family of proteins that function as epigenetic modifiers and transcriptional regulators (Chittock et al. 2017). Two key complexes are polycomb repressive complex 1 PRC1, an E3 ubiquitin ligase that monoubiquitinate lysine 119 of histone H2A (H2AK119ub1), and polycomb repressive complex 2 (PRC2), which functions as a methyltransferases that targets histone H3 lysine 27 for mono-, di-and trimethylation (H3K27me1, 2, 3). Amongst genes that branched between PGCs and soma, we noted a particularly strong overrepresentation for targets of PRC1 and PRC2. Indeed, PRC-related terms were amongst the most frequently enriched terms when comparing branched genes with ChEA/Encode databased (Supplementary File 3). This included enrichment for targets of core PRC2 components SUZ12 (p<1×10^-23^, Enrichr ChEA) and EZH2 (p<1×10^-10^, Enrichr ChEA), and proteins known to bind or co-localise with PRC2, including JARID1A (p<1×10^-51^, Enrichr ChEA), JARID1B (p<1×10^-73^), JARID2 (p<1×10^-8^; ChEA), and KDM6A (5×10^-21^, Enrichr ChEA). We also saw enrichment for PRC1 components RING1B (5×10^-10^, Enrichr ChEA) and RNF2 (5×10^-10^; Enrichr ChEA), as well as non-canonical components such as YY1 (1×10^-58^, Enrichr ENCODE) and MAX (p<1e10^-52^, Enrichr ENCODE). The expression patterns of PRC1/PRC2 components following pseudotime ordering revealed highly dynamic regulation (Supplementary Figure 6). K-means clustering of gene bodies of early-branching genes (*t_b_* < 0.6) by RING1B occupancy and ubiquitination level in hESCs (GSE104690) allowed identification of putative targets of PRC1, which included the key PGC specifier SOX17 (Supplementary Figure 7 and 8).

Together these results suggest PRC1/2 may play an important role in PGC development. To establish a possible role of PRC1/2 on PGC specification, we used small molecule inhibitors to target the enzymatic activity of core components of PRC1 and PRC2 using an *in vitro* model of PGC specification. Our results reveal that inhibition of PRC1 RING1B/BMI1-dependent ubiquitination using PRT 4165, reduced PGC specification in a dose-dependent manner (Supplementary Figure 9). Previous studies suggest that the small molecule inhibitor DETA/NONOate could be used to suppress expression of YY1 (Hongo et al. 2005). Treatment with DETA/NONOate reduced PGCLC efficiency (Supplementary Figure 10), however, DETA/NONOate is also known to activate the NO signalling pathway, and we could not rule out a YY1-independent mode of action. Finally, a preliminary inhibition of PRC2 EZH2-mediated histone methylation via DZNep further suggested reduced efficiency of PGCLC specification (Supplementary Figure 11). These results highlight the ability of pseudotime models to identify targets for intervention that result in a phenotype.

#### 3.2 Differences between inferred in vivo and in vitro dynamics of human development

Current knowledge of human germ cell specification is largely informed by *in vitro* PGCLC derivation from hESCs. While it is challenging to directly validate these findings *in vivo*, we reasoned that we could highlight the most relevant processes by comparing the inferred dynamics of PGC induction in early embryos with pseudotemporally ordered hESCs. In particular, the ability to derive human PGC-like cells (PGCLCs) from hESCs *in vitro* prompted us to regard the hESC to hPGCLC transition as a recombination process. Comparison of the PGCLC transcriptome with that of *in vivo* PGCs suggests that PGCLCs closely resemble pre-migratory PGCs (Irie et al. 2015), and we therefore considered recombination to occur between hESC and the earliest available PGC dataset (Wk4). Thus, the comparison of hESC and hPGC dynamics represents a branch-recombinant process, whereby specified hPGCs and hESCs branch from a common precursor (around blastocyst stage), with hESC dynamics allowed to recombine with W4 hPGCs upon exposure to appropriate stimuli.

We first identified the subset of genes that showed divergent behaviour between ESCs and *in vivo* datasets. This revealed 1,331 branching genes which were up-regulated in hESCs compared to *in vivo*. The timing of divergence between inferred *in vivo* and *in vitro* dynamics is shown in Supplementary Figure 4(c). GO analysis of these groups identified several enriched terms, including those relating to cell adhesion, response to hormones, and, importantly, response to BMP4 and WNT, as well as terms relating to germ cell development and meiosis (see Supplementary File 3). We next attempted to identify perturbations that could potentially drive hESCs back towards hPGC identity i.e., for genes up-regulated in hESCs versus hPGCs, we searched for perturbations that down-regulated those genes.

Amongst the most highly enriched signatures was the overexpression of *SOX17* (p<5×10^-9^, Enrichr SG). D*e novo* motif analysis using DREME (Bailey 2011) on 1kb windows upstream of the TSS identified two motifs (p<1×10^-17^, p<5×10^-7^) resembling that of human *PRDM1* (p<0.01, p<5×10^-3^, TOMTOM, Gupta et al. (2007)), and genes were enriched against DE genes in bulk RNA-seq data from *PRDM1*-overexpressing cells (Kobayashi et al. 2017) showed a significant overlap (p<5×10^-24^, hypergeometric test).

A strong signature was also noted for *KIT* (p<1×10^-3^, Enrichr SG), potentially highlighting an important role of its ligand, SCF, in PGC cell fate. SCF has been implicated in PGC proliferation and migration (Hoyer, Byskov, and Mollgard 2005; Mollgard et al. 2010) and is added to the *in vitro* culture medium used to derive PGCLCs from hESCs (Irie et al. 2015). Although *KITLG* is not expressed in hPGCs or hPGCLCs (Tang et al. 2015; Irie et al. 2015), the expression of *KIT* is significantly up-regulated in hPGCs, and SCF (encoded by *KITLG*) is potentially secreted from adjacent cells. Indeed SCF is expressed in Schwann cells (Mollgard et al. 2010), as well as Sertoli and Leydig cells (Sandlow et al. 1996), and we note that a *KITLG* positive subpopulation appears to exist in the somatic cells, around the time of PGC migration and gonad colonisation.

#### 3.2.1 Early branching identifies key regulators of competence for PGC fate

Human ESCs in conventional cultures (KSR-based medium supplemented with FGF2) have low potential to form germ cells, but acquire competency for PGCLC fate in the presence of TGFβ, LIF, FGF2 and the inhibitors of GSK3β (CH), p38 (SB), JNK (SP) and MEK (PD03) kinases (4i medium; Gafni et al. (2013); Irie et al. (2015)). 4i hESCs self-renew in a competent state and can form high numbers of PGCLCs when exposed to BMP2/4 and supporting cytokines in embryoid body (EB) cultures (Irie et al. 2015). The mechanisms underlying such dramatic change in developmental potential of 4i (competent hESCs) versus conventional hESCs remain to be elucidated.

We decided to focus on signalling that may be involved in conferring PGC competence to hESCs. For this, we first compared the genes up-regulated in hESCs to the inferred *in vivo* dynamics for PGCs. This revealed enrichment for the downregulation of BRAF (p<1×10^-8^, Enrichr SG) and FGF2 (p<1×10^-12^, Enrichr Ligand), two upstream regulators of MEK signalling. Crucially, FGF2-responsive genes were found to branch very early *in vivo* (soma vs PGCs) compared to overall branch times (p<5×10^-6^, KS-test) and was the topmost enriched ligand perturbation identified from comparisons of ESCs towards PGC fate (Suppelmentary File 5). This branching occurred concurrent with, or prior to the BMP4/SOX17-responsive genes (p<0.05, FGF2 vs BMP4; NS, FGF2 vs SOX17, see Figure 2(h)), suggesting that FGF2 may function earlier than key specifiers of PGC fate, with potential roles in conferring competency for PGCLC fate. While FGF2 is present in both conventional and 4i conditions, 4i cells are maintained in the presence of an inhibitor of MEK, an effector of FGF receptor (FGFR) signalling. It is therefore possible that partially negating the effect of FGF signalling via the inhibition of downstream pathways of FGFR, might contribute to PGC competence acquisition by pluripotent cells.

Comparison of branching between soma and PGCs identified early signatures for a number of kinase perturbations that mimic the use of the inhibitors in the 4i medium sustaining competent hESCs. These included: (i) knockdown of GSK3B (p<5×10^-6^, Enrichr Kinase; p<0.05, KS-test) (CH is a GSK3β inhibitor); (ii) knockdown of MAP2K1 (p<1×10^-3^, Enrichr; p<0.05, KS-test), a component of the MAP kinase signal transduction pathway upstream of MEK signalling (PD03 is a MEK inhibitor); and (iii) MAPK14 (p<5×10^-5^, Enrichr; p<0.01, KS-test), a regulator of p38 (SB is a p38 inhibitor). Branch times for genes associated with these perturbations were suggestive of early roles. Of note, we did not identify statistically significant overlaps with pathways downstream of JNK kinases, inhibited by the fourth inhibitor in 4i hESCs (SP).

To experimentally test the involvement of these signalling pathways in PGC competence, we compared PGCLC induction efficiencies from hESCs cultured in either complete competent (4i) medium (control) or lacking one of the 4 inhibitors: (i) “4i-CH” (4i without GSK3B inhibitor); (ii) “4i-PD03” (4i without MEK inhibitor); (iii) “4i-SB” (4i without p38 inhibitor), and (iv) “4i-SP” (4i without JNK inhibitor). hESCs were cultured in these alternate media for at least one passage and collected for standard PGCLC differentiation (as in Irie et al. (2015)). PGCLC induction efficiency was then quantified by flow cytometry (Supplementary Figure 12). This was facilitated by the use of PGC-specific knockin reporter cell line NANOS3-tdTomato (N3tdTom; Kobayashi et al. (2017)) and staining for PGC surface markers AP and CD38 (Irie et al. 2015).

This identified that while removal of CH (GSK3βi) and SP (JNKi) from 4i did not significantly change PGCLC competence, withdrawal of PD03 (MEKi) and SB (p38i) strongly reduced the propensity of hESCs to form PGCLCs (Figure 3(a)), consistent with a predicted role for MEK and p38 but not JNK signalling in conferring PGC competence. Considering the crucial role of WNT signalling for PGC competence in mice (Ohinata et al. 2009; Aramaki et al. 2013), it was surprising that CH (WNT agonist, Ying et al. (2008)) withdrawal did not impact on PGCLC competence. We hypothesized that 4i hESCs might produce WNT in an autocrine fashion. Indeed, the number of PGCLCs generated from hESCs grown with WNT pathway inhibitor IWR-1 (IWR, Chen et al. (2009)) either in the presence or the absence of CH, was reduced compared to the control (Figure 3(b)).

Notably, “4i-CH” and “4i-SB” cultures deteriorated at later (>4) passages, reflecting the importance of these inhibitors to sustain self-renewal of 4i hESCs (Gafni et al. 2013). Low competency observed upon withdrawal of SB (p38i) was likely caused by the induction of trophectoderm-like differentiation, as judged by the expression of lineage markers, *CDX2* and *HAND1* (Supplementary Figure 13).

The most drastic effect on PGCLC competence was observed upon withdrawal of MEK inhibitor, PD03, with a strong reduction in the induction of PGCLC (Figure 4(b)), although unlike “4i-CH” or “4i-SB”, “4i-PD03” cells could be maintained for many passages (>20). Importantly, the defect of “4i-PD03” hESCs to differentiate to PGCLCs could be rescued (within one passage) if PD03 was reintroduced; Figure 4(c)).

Since our analysis revealed early branching of an upstream component of this signaling pathway, namely FGF2 (see Supplementary File 3), we asked if inhibition of FGF receptor (FGFR) could mimic the effect of MEK inhibition. To address this, we transferred the cells cultured in “4i-PD03” to “4i-PD03+PD17” (4i where MEK inhibitor PD0325901 was substituted for FGFR inhibitor PD173074). Not only did FGFR inhibition rescue the differentiation defect of “4i-PD03” cells, but it also markedly enhanced PGCLC competence compared to the 4i control (in the presence of high PD17 concentrations; Figure 4(d)). However, these cells showed decreased viability and collapsed after 2-3 passages. Importantly, FGFRi showed a dose-dependent positive effect on competence (Figure 4(d)) and cells grown with lower PD17 concentration could be maintained longer. Together, these data show that, in agreement with our computational prediction, FGF-MEK pathway is a negative regulator of human PGCLC competence; blocking this cascade by FGFRi or MEKi promotes PGCLC competence in hESCs.

Next, we explored the reason for enhanced competence of FGFRi-treated versus MEKi-treated cells. Major downstream components of FGFR signalling are RAS-MEK-ERK, PI3K-AKT, and JAK-STAT (Lanner and Rossant 2010). Indeed, the comparison of dynamics of PGC development with somatic cells identified strong enrichment for these three pathways.

Thus, we saw enrichment of appropriate GO/KEGG terms, and transcriptional signatures of EGFR (p<1×10^-19^, Enrichr Kinase; p<0.0005 KS-test) and Platelet Derived Growth Factor Receptors Alpha (*PDGFRA*, p<1×10^-11^, Enrichr Kinase; p<0.005 KS-test) and Beta (*PDGFRB*, p<5×10^-7^, Enrichr Kinase; p<0.001 KS-test), implicated in the activation of ERK, AKT, and STAT1/3/5 signaling pathways, as well as *MET* (p<5×10^-10^, Enrichr Kinase; p<0.01 KS-test), which functions upstream of RAS-ERK and PI3K-AKT signalling. We also noted a strong signature for the down-regulation of *JAK1/2* (p<5×10^-14^, Enrichr Kinase; p<0.05 KS-test), and an enrichment for PI3K signalling, which included transcriptional signatures of *AKT1* (p<1×10^-9^, Enrichr Kinase; p<0.05 KS-test), *CSF1R* (p<5×10^-5^, Enrichr Kinase; p<0.01 KS-test), the regulatory subunit of PI3K leading to activation of AKT1, and *NTRK3* (p<5×10^-5^, Enrichr; p<0.05 KS-test), which also activates PI3K-AKT signalling. PI3K/AKT pathway was of particular interest, since it is activated downstream of insulin signalling, which was one of the most statistically enriched ligand perturbations from the Enrichr database (p<1×10^-7^, Enrichr ligand; p<0.01 KS-test) promoting hESC to hPGC transcriptome transition.

We therefore hypothesized that simultaneous inhibition of MEK and other pathways downstream of FGFR could mimic the effect of FGFRi and increase PGCLC competence. To test this, we supplemented complete 4i cultures with either LY (*LY294002*, PI3K inhibitor) or CAS (CAS457081-03-7, JAK inhibitor). This revealed a trend for enhanced PGCLC competence of “4i+LY” hESCs (Figure 5(a)). Importantly, “4i+LY” cells did not exhibit increased cell death as seen with FGFRi; instead they proliferated at rates similar to 4i hESCs and formed homogeneous dome-shaped colonies. Furthermore, “4i-PD03+LY” cells, especially at later passages exhibited competence comparable to control 4i hESCs (Figure 5(a)), pointing to a synergistic action of MEK/ERK and PI3K/AKT signalling pathways. We also observed a positive effect of JAK pathway inhibition on PGCLC competence (Figure 5(b)). This is in line with the enhanced competence of hESCs cultured in 4i without LIF, an agonist of JAK-STAT (Figure 5(c)). Interestingly, differentiation of 4i+CAS” hESCs yielded more CD38+ cells (Figure 5(c)), which represent more mature PGCLCs (Irie et al. 2015). Of note, longer culture of hESCs with JAK inhibitor changed colony morphology and subsequently reduced competence (Figure 5(b)). These cultures could not be maintained, highlighting differential requirements for JAK/STAT signalling in 4i versus conventional hESCs (Gafni et al. 2013; Onishi and Zandstra 2015). It is therefore possible to speculate that the observed effect of FGFR inhibition could be explained by: (i) enhanced competence due to PI3K pathway inhibition and transient JAK/STAT inhibition; (ii) decreased viability and loss of pluripotency due to JAK/STAT inhibition. The relationship between these signalling pathways and their contribution to pluripotency maintenance and PGC competence acquisition warrant further investigation. Together, these data show that B-RGPs could predict the relevance of specific signalling pathways to PGC competence even in the absence of all relevant data points (single-cell transcriptomes of 4i (competent) hESCs and PGCLCs).

**Figure 5.**
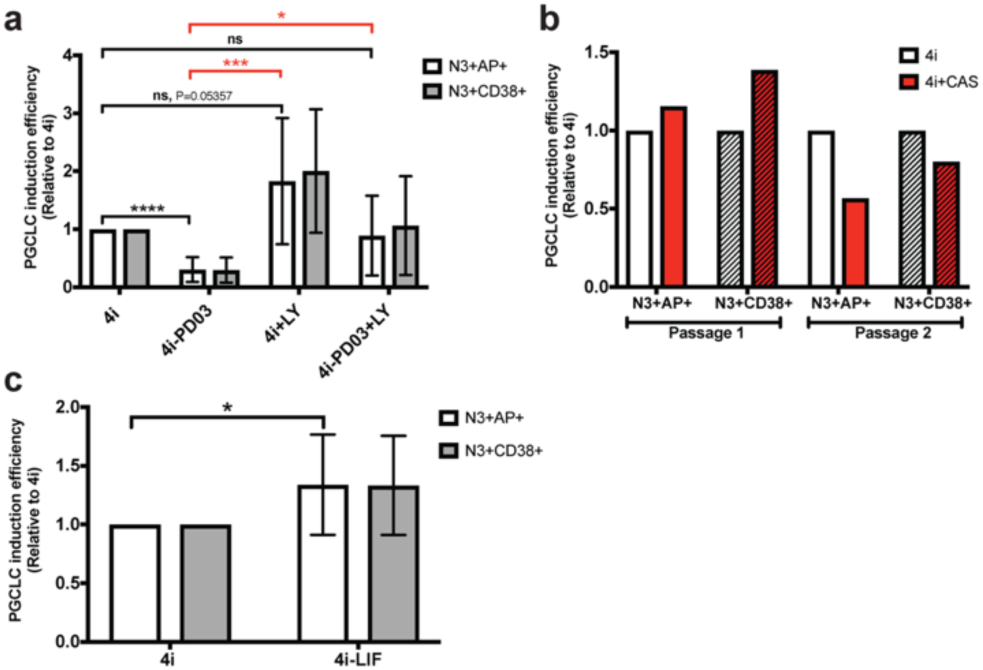
PI3K and LIF/JAK/STAT pathways negatively regulate PGCLC competence. **(a)** PI3K inhibition promotes PGCLC competence. Quantification of PGCLC induction efficiency from cells grown in indicated conditions relative to 4i hESCs. Data are shown as mean ± SD of 10 independent experiments. * p ≤ 0.05, *** p ≤ 0.001, **** p ≤ 0.0001, ns: not significant (p > 0.05), Holm-Sidak t-test (on relative frequency of N3+AP+ cells). Red lines and asterisks refer to comparison of “4i-PD03” to other conditions. **(b)** Short exposure (1 passage) of hESCs to JAK inhibitor CAS increases PGCLC competence to form mature CD38+ PGCLCs Quantification of PGCLC induction efficiency from cells grown in 4i+CAS relative to 4i hESCs. Longer culture with CAS changed colony morphology and decreased competence. **(c)** LIF withdrawal from 4i hESCs promoted PGCLC competence, although the magnitude of the effect was mild. Data are shown as mean ± SD of 10 independent experiments. * p≤ 0.05, Holm-Sidak t-test (on relative frequency of N3+AP+ cells).

Finally, we set out to test if in addition to promoting competence, blocking MEK signalling could also enhance PGCLC specification. To this end, we supplemented the differentiation medium with MEK inhibitor PD03 or induced a genetic construct for dominant-negative ERK expression (DN-ERK) at the onset of differentiation (Figure 6(a)). Intriguingly, both perturbations reduced PGCLC numbers compared to controls (Figure 6(b, c)). ERK activation during PGCLC induction is likely triggered by the supplementation of the differentiation medium with EGF (Irie *et al*., 2015), which activates similar intracellular pathways to FGF ligands (Schlessinger 2004). Of note, inhibition of PI3K throughout differentiation did not abrogate PGCLC specification, suggesting that, unlike MEK, PI3K signalling is dispensable for PGCLC induction (Supplementary Figure 15). Altogether, this shows that the timing of FGFR signalling is crucial for context-specific cell fate decisions and our methodology allows predicting relevant changes in signalling and gene expression in a time-resolved manner.

**Figure 6.**
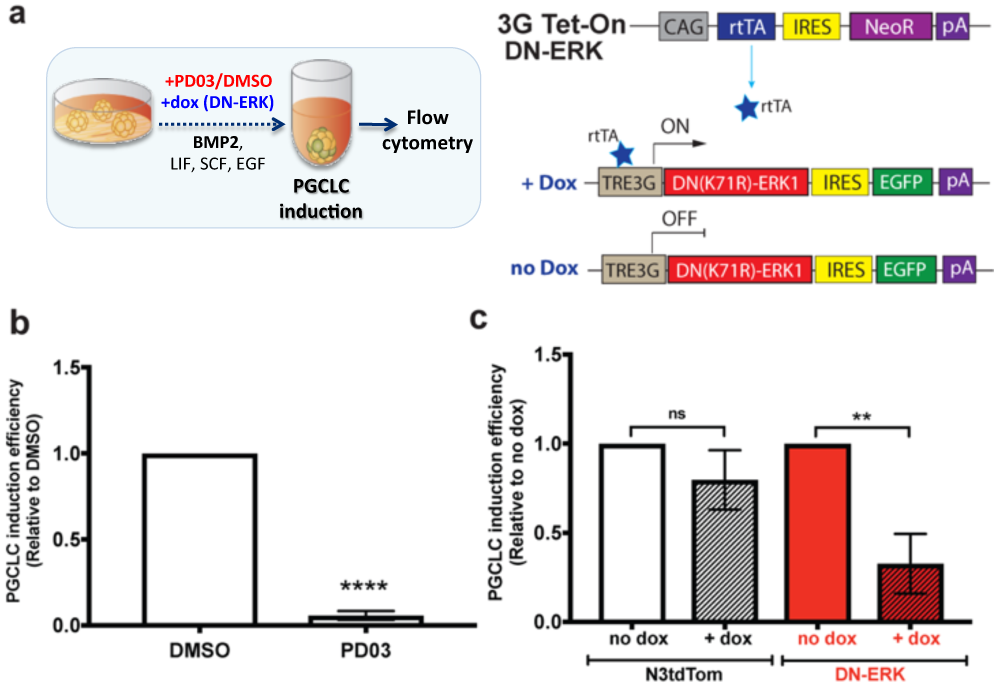
MEK-ERK inhibition during PGCLC induction interferes with PGCLC specification. **(a)** Workflow of the experiments and the scheme of the construct for DN-ERK transgene expression. Either PD03 or DMSO (vehicle) was added to the differentiation medium Alternatively, to induce DN-ERK, PGCLC medium was supplemented with doxycycline (dox) **(b)** Supplementation of the differentiation medium with PD03 drastically decreases PGCLC specification efficiency. Data (relative frequency of live N3+AP+ cells) are shown as mean ± SD of 4 independent experiments. **** P ≤ 0.0001, Holm-Sidak t-test. **(c)** DN-ERK expression during PGCLC differentiation reduces the efficiency of PGCLC specification. Data (relative frequency of live N3+AP+ cells) are shown as mean ± SD of 3 independent experiments using 2 DN-ERK-expressing clones. ** p ≤ 0.01, Holm-Sidak t-test.

## 3 Discussion

Transcriptional branching and recombination are frequently encountered in developmental biology. Single-cell transcriptomics has emerged as tool for investigating the nature of these bifurcations, but requires cells first be computationally ordered along pseudo-developmental trajectories. Increasingly, scRNA-seq datasets are generated as part of more structured experiments, including finely resolved whole-embryo developmental time series (Briggs 2018). Unfortunately, most pseudotime approaches cannot, or do not, leverage the additional information. Here we have developed a probabilistic model capable of inferring a posterior distribution over pseudotimes, that also utilises prior information about capture. Our approach encodes an explicit model of a bifurcations occurring at the level of individual marker genes, providing a more interpretable result than standard dimensionality reduction.

Although our model was more accurate than other approaches for time series data, this increased accuracy comes at a significantly increase in computational cost. However, concurrent with our study, Boukouvalas, Hensman, and Rattray (2018) have built on the earlier work of Yang et al. (2016) to develop Branching Gaussian processes, a probabilistic framework for the inference of bifurcations in single cell transcriptional datasets. Unlike our approach, which utilised MCMC to deal with unlabelled datasets, these branching GPLVM focused on efficient, scalable variational approximations for a two-branch system, implemented using GPflow (Matthews et al. 2016), and were therefore applicable for much larger datasets. By focusing on early branching events they identified key regulators of haematopoietic differentiation which, together with our own work, and other GPLVM approaches (Ahmed, Rattray, and Boukouvalas 2018), demonstrates the usefulness of probabilistic approaches to pseudotime.

Having established the advantages of B-RGPLVMs over other approaches for time series scRNA-seq data, we used our approach to investigate the dynamics of cell fate decision of human primordial germ cells (hPGCs). We correctly identified key early regulators of PGC fate, most notably *SOX17*, a classic endoderm marker gene that was only recently shown to play a prominent role in human PGC specification (Irie et al. 2015; Tang et al. 2015). Furthermore, by jointly leveraging *in vivo* and *in vitro* datasets, we could suggest the importance of correct suppression of *SOX2* and concomitant up-regulation of *SOX17* in hPGC lineage. Interestingly, mouse PGCs do not require *Sox17* and instead express *Sox2*. Many more unexplored differences exist between mouse and human PGC transcriptomes (Tang et al. 2015), and an integrative analysis of mouse and human datasets could prove useful in identifying such divergent regulators.

By looking at the earliest branching events, we identified putative regulators that may play a role prior to PGC specification, in the acquisition of PGC competence. Here, we identified possible roles for FGFR signalling and its downstream branches (MEK/ERK, PI3K/AKT and JAK/STAT) in conferring human PGC competence. Importantly, these observations were validated experimentally using an *in vitro* system for derivation of PGC-like cells (PGCLCs) from hESCs. Indeed, blocking FGFR or its downstream effectors (MEK, PI3K and JAK) by specific inhibitors enhanced the competency of hESCs to form PGCLCs.

We also noted that branched genes were highly enriched for PRC1 and PRC2 binding. Preliminary small molecule inhibition of the enzymatic components of RING1B and EZH2 resulted in reduced ability of competent ESCs to form PGCLCs. PRC1/2 components showed highly dynamic pseudotime trajectories, both around the time of specification and in post-migratory PGCs, suggesting that PRC1/2 components such as MAX may have further important roles in later hPGC development, consistent with studies in mouse models (Yokobayashi et al. 2013; Suzuki et al. 2016; Endoh et al. 2017).

Whilst we could correctly identify several events in the acquisition of competency and specification, key intermediate cell types were absent from both our *in vivo* and *in vitro* datasets. Notably, single cell RNA-seq for competent (4i) hESCs and PGCLCs were unavailable, although bulk RNA-seq measurements exist (Irie et al. 2015). A key future development will therefore aim to combine the use of single-cell and bulk RNA-seq data in a principled way. Since bulk measurements represent population averages, B-RGPs would be ideally suited to this purpose, due to the ability of GPs to incorporate integral observations (Rasmussen and Williams 2006). Likewise, as GPs can naturally incorporate derivative observations, GPLVMs provide an ideal framework for leveraging other useful information such as RNA velocity (La Manno et al. 2018).

Another informative approach would require generation of single cell transcriptomic profiling of intermediate cell types, providing a higher temporal resolution of the intermediate events that lead to the acquisition of competence and specification of hPGCs. The emergence of other *in vitro* models for human PGCLC derivation (Kobayashi et al. 2017), as well as *in vitro* models of embryogenesis (Harrison et al. 2017; Sozen et al. 2018; Beccari et al. 2018; Deglincerti et al. 2016; Martyn et al. 2018), provides further opportunities for dissecting these causal regulations; by identifying differences and similarities in the dynamics of branching between different *in vitro* models, we could separate out the underlying biological mechanisms from culture-induced adaptations.

Finally, the transition from pluripotency to PGC competence and ultimately to PGCs can be reversed later in development upon germ cell tumour formation, exemplifying a recombination event. Seminoma and embryonal carcinoma are two types of germ cell tumours that share similarities with PGCs and ESC, respectively (Surani 2015), which can be distinguished using the markers identified from *in vitro* human PGCLC specification; seminomas and PGCs express CD38 and SOX17, while embryonal carcinomas and hESCs express SOX2 and CD30 (Irie et al. 2015). This underscores the importance of interrogating the transcriptional and epigenetic control of human germ cell fate and its specification from pluripotent progenitors. Ultimately, the use of B-RGPLVMs on transcriptomics data from tumour cells could shed light on the sequence of events that lead to their formation, identify cancer markers, and guide therapeutic interventions.

## Acknowledgements

We thank Ufuk Günesdogan and Naoko Irie for critical reading of the manuscript, Toshihiro Kobayashi for the provision of figures and advice, and the Surani lab for useful insights. We thank Charles Bradshaw for help with high performance computing, and the Gurdon Institute core facilities for continued support. Finally, we thank Magnus Rattray, Jing Yang, James Hensman, and Alexis Boukouvalas for useful insights and discussion about statistical branching models.

## 1 Branch-recombinant Gaussian process latent variable models (B-RGPLVMs)

The ability to leverage capture time and other important prior information into pseudotime algorithms, as well as the ability to quantify uncertainty in the pseudotemporal ordering, is particularly desirable given the inherent limitations of pseudotime approaches (see discussion in Campbell and Yau (2015); Weinreb et al. (2017)). Unfortunately, most approaches for pseudotime ordering fail to incorporate such information, a significant omission given the trend towards increasingly structured (finely resolved time-series) scRNA-seq datasets.

Bayesian approaches represent an ideal framework for leveraging prior information, and previous studies by Reid and Wernisch (2016) have demonstrated how capture time can be incorporated into pseudotime algorithms using Bayesian approaches based on Gaussian process latent variable models (GPLVMs; Lawrence (2003,2005)). Within these models, it is assumed that there are *M* cells measured at one of *T* < *M* distinct known capture times, ***t***_*c*_ = (*t*_1_, …, *t_M_*,), and the aim is to infer a corresponding set of pseudotimes, ***t***_*p*._ = (*τ*_1_, …, *τ_M_*,), such that the gene expression profiles vary smoothly over pseudotime in a way that reflects the general developmental trajectory. The pseudotimes, **t_p_**, are assigned a prior distribution that is Gaussian distributed conditional on the capture time, **t**_*c*_, **t**_*p*._|**t**_*c*_ ~𝒩 (**t**_*c*_, *σ*^2^ 𝕀). Within the GPLVM the expression profile *f_i_* of each gene *i* is assigned an independent Gaussian process prior, 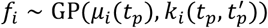, where *μ_i_*(*t_p_*) denotes the mean function and 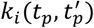 the covariance function. Different choices of covariance function encode different prior distributions over the expression profiles of the individual genes, and can be used to encode a vast range of dynamic behaviour. Reid and Wernisch (2016) assume a Matérn covariance function:

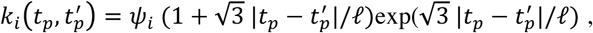

where *ℓ* is a global length-scale and *ψ_i_* are gene-specific scaling factors. The expression data for each gene *i* over the observed cells is modelled as a noisy version of their underlying expression profile, 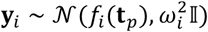, where *ω_i_* represents gene-specific noise levels. As the Gaussian process prior is conjugate to this likelihood, *f_i_* can be directly marginalised:

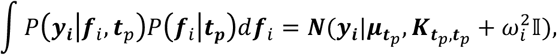

where 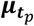 represents a vector of the mean function evaluated at the pseudotimes times ***t***_*p*_, and 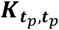 the covariance matrix. Finally, the composite likelihood for pseudotimes can be written as:

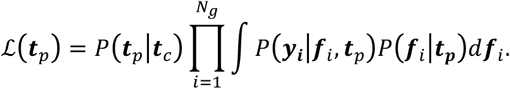

In general, it is not computationally feasible to evaluate this product over all genes, and a representative set of genes must be selected instead. This might include taking known marker genes, which has previously been shown to accurately order cells in other approaches (Campbell and Yau 2018), or using the top most variable genes (Reid and Wernisch 2016). Until recently, no explicit GP treatment for branching processes existed, and these kinds of models were thus limited to single developmental trajectories (Reid and Wernisch 2016). However, recent studies by Yang *et al.* (2016) have derived explicit covariance functions for a two-branch process allowing pseudotime approaches for two-branch systems (Boukouvalas, Hensman, and Rattray 2018), and subsequent studies have demonstrated how compositions of covariance functions can be used to define branching processes of arbitrary complexity (Penfold et al. 2018). Within this paper, we will consider processes with one or two bifurcations. A two-branch system with observations at ***t*** = (*t*_1_, …, *t_M_*), and branch-labels, ***z*** = (*z*_1_, …, *z_M_*), *z_i_* ∈ [1,2], can be described by the following correlated processes:

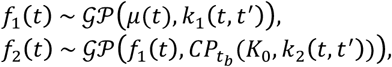

Where *μ*(*t*) represents the mean function for the base process, *K*_0_= *K*_0_ (*t,t′*) denotes a zero-kernel, and (*k*_1_, *k*_2_) denotes a change-point kernel (Lloyd et al. 2014), defined as:

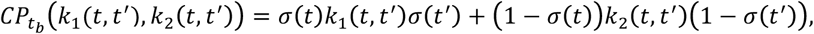

Where 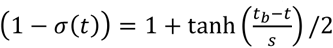. In this system the hyperparameter, *t_b_*, controls the time at which the second trajectory diverges from the basal process. A three-branch process with observations at identical times as above and branch labels ***z***= (*z*_1_,…,*z_M_*), *z_i_* ∈ [1,2,3], can be defined via the following set of correlated processes:

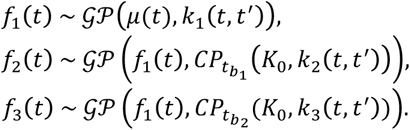

For this system, cells corresponding to label 1 can be interpreted as the basal developmental process, from which two developmental programs independently diverge at times 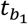 and 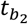. Finally, we can consider a three-branch system, where cells correspond to branch 2 diverging at time 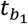, and cells corresponding to branch 3 diverging at 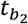 and later reconverging at time 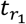 This system is described by the following coupled processes:

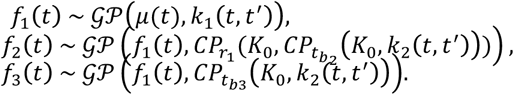

As with the standard GPLVM, each gene can be assigned an independent B-RGP prior over the pseudotimes, and the aim is to order cells over a smooth, bifurcating process. The composite likelihood for the B-RGPLVM is:

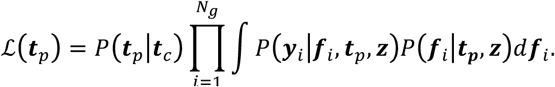

Where ***y***_*i*_ is the vetctor of gene expression for the *i*th gene. While inference of the posterior distribution over pseudotimes is analytically intractable, we may readily sample from it using Markov chain Monte Carlo, with pseudotimes updated via a Metropolis step, hyperparameters sampled via hybrid Monte Carlo and, where necessary branch labels updated via a Gibbs step. We can apply a perturbation to the pseudotime 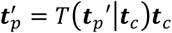 which is accepted with probability *P* (*accept*) = min(1, *A*), where:

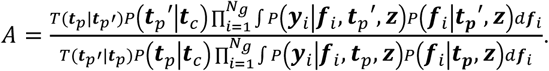

Other updates can be applied, for example the swapping of two randomly selected cells, and more principled approaches to such permutation-based updates for pseudotime are developed by Strauss, Reid, and Wernisch (2018). Finally, if we are dealing with situations where cell fate is uncertain, we can Gibbs sample the branch label for cell *i*:

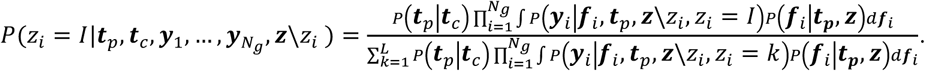

where *L* represents the number of cell types.

### 1.1. Benchmarking B-RGPLVM

To assess our ability to pseudo-temporally order data over a branching process using B-RGPLVMs, we first benchmark using existing microarray time series data measuring the *Arabidopsis thaliana* transcriptional response to the necrotrophic bacteria, *Botrytis cinerea* (Windram et al. (2012); GEO GSE39597). This dataset consists of two time-series: (i) a control time series, detailing changes in gene expression in *Arabidopsis* over a 48-hour period at two-hour intervals; and (ii) a time-matched infection series, in which Arabidopsis has been inoculated with *B. cinerea*. The infection time series has previously been used to benchmark GPLVMs for pseudo-time ordering (Reid and Wernisch 2016). As outlined by Reid and Wernisch (2016), we first grouped the individual measurements into four groups containing six consecutive time points, artificially reducing the temporal resolution of each time series dataset from 24 time points to 4. We then attempted to recapture the correct ordering of cells using an increasing number of randomly selected genes within the B-RGPLVM. Here we selected 10, 20, 40, and 80 genes at random from the set of 150 previously used by Reid and Wernisch (2016). By doing so, we aimed to identify how the accuracy of the model changed with respect to increased number of gene observations, and thus empirically identify the number of genes required for accurate pseudotime ordering. We performed 5 different randomly initialised runs, with 10,000 samples in the MCMC chain, discarding the first 5,000 steps for burn-in. Run time was in the region of 10 hours, although this could be decreased using MATALBs parallel processing toolbox. Note that for each independently initialised run, a separate random selection of genes was chosen to gauge variability in the inferred ordering with respect to different genes.

Since the time series were generated using bulk microarrays, with measurements based on populations of cells, we expected a smoothly varying process, and thus for the covariance function we used a two-component branching covariance function composed of squared-exponential covariance functions.

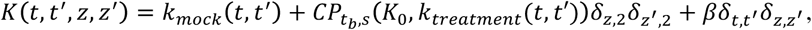

where 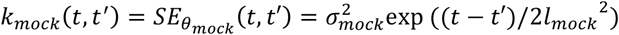 and 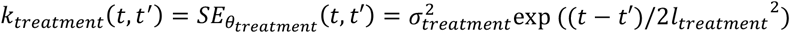. The hyperparameters in the model therefore corresponded to the length-scale and process variance of the base process (the control data), the length-scale and process variance of the perturbed process (the infected data), and the change point time (branch time) and branch rate for the change-point kernel. Hyperparameters were initially fitted using maximum likelihood estimates based on the low-resolution capture times, θ ← arg_*θ*_ max 𝓛(*θ*|***t***_*p*_ = ***t***_*c*_, ***y***). Length-scale, process-variance, and noise hyperparameters were then fixed for the remainder of the inference. For the change-point time hyperparameter, *t_b_*, we assumed a univariate smoothed box prior distribution with linear decay in the log domain, *P*(*t_b_*|*a,b*) = *σ*(*η*(*t_b_* – *a*))(1 – *σ*(*η*(*t_b_* – *b*))), where *σ*(*z*) = 1/(1 + exp (–z)),and *a* = 0, *b* = 33, and *η* = 33.

In Supplementary Figure 1, we plot the inferred pseudotime ordering for control and infection datasets versus the true ordering of data as the number of genes used within the GPLVM increased, with *N* ∈ (10,20,40,80). We note that, in general, the accuracy of the models appears to increase as the number of genes increases. For the case *N* = 40, 4 out of 5 replicates were nearly perfectly ordered for both the infected and control time series, with a mean correlation of >0.97 and >0.98 respectively over the five runs (for N=40). No obvious improvement was seen when increasing the number of marker genes to 80 (Supplementary Figure 1(c)). Consequently, when pseudo-temporally ordering data, we used >40 genes.

For comparison purposes, we also performed pseudotime using the full list of 150 marker genes previously used by Reid and Wernisch (2016) as taken from the main text of Windram et al. (2012). Here we used two approaches:

- Monocle2 (Qiu et al. 2017). We first ran Moncle2 using the combined control and treated datasets. Subsequently we ran it on the control and treated datasets separately. The later analysis showed higher correlation between inferred pseudotime and measurement time and are the results reported here.
- TSCAN (Ji and Ji 2016). We used the online TSCAN app (https://zhiji.shinyapps.io/TSCAN) on the control and treated datasets separately. As the data was already log normalised we applied no normalisation. We used PCA for dimensionality reduction, with number of components set using the ‘Automatically select optimal dimension for PCA’ option. Finally the number of clusters was selected with the ‘Use optimal cluster number’ option, and pseudotimes exported as csv files.

In supplementary Figure 2 we plot the measurement time versus inferred pseudotime for Moncle2 and TSCAN respectively for the control and infected datasets. For TSCAN we saw Pearson correlation of 0.74 and 0.81 respectively in the control and infected datasets, whilst for Monocle2 we found a correlation of 0.85 and 0.83 respectively. For the BRGPLVM we found a mean correlation of 0.97 and 0.99 respectively over five runs (for N=40).

## 2 PSEUDOTEMPORAL ORDERING OVER DEVELOPMENTAL TRAJECTORIES

Single cells from pre-implantation embryos, PGCs, somatic cells and ESCs (Guo et al. 2015; Yan et al. 2013) were initially pseudotemporally ordered over a two-component branching process using 44 marker genes. For each of the 44 genes, the trajectory for specification of PGCs was chosen to represent the base process, due to this class having the most data points, with developmental trajectories for soma representing the branch process. Pre-blastocyst stage cells were randomly assigned to either branch with equal probability, whilst branch labels for blastocyst stage cells and ESCs were inferred within the algorithm. We performed five randomly initialised runs, using 30,000 steps in the MCMC chain, in some cases taking >48 hours using a single CPU, although key bottlenecks are embarrassingly parallel, and runtime could be reduced dramatically using the MATLAB parallel computing toolbox. The order of cells at step 30,000 appeared to show good overall correlation across these five runs, with mean correlation coefficient, 〈*R*〉 = 0.9768 ± 0.003.

For comparison purposes we also pseudotemporally ordered cells using a variety of other pseudotime methods, including GrandPrix (Ahmed, Rattray, and Boukouvalas 2018), Monocle2 (Qiu et al. 2017), SLICER (Welch, Hartemink, and Prins 2016), and SCUBA (Marco et al. 2014), TSCAN (Ji and Ji 2016), Wishbone (Setty et al. 2016).

- GrandPrix (Ahmed, Rattray, and Boukouvalas 2018). We ran GrandPrix using data from the soma and PGC lineages, capture times were first scaled lie in the interval [0, 1]. We used two latent dimensions, with cells assigned Gaussian priors based on capture time along the first latent dimension and standard deviation of 0.05. Ordering was chosen based on the position along latent dimension 1.
- Moncole2 (Qiu et al. 2017). Here we ran Monocle2 several times. Initially we included all *in vivo* cells and attempted to capture the bifurcations between soma and PGCs. Within the algorithm, we used SOX17 and WT1 as marker genes for terminal fates of PGC and soma, respectively. In the second instance, we separated out the data, first combining pre-implantation data with PGCs, to infer the PGC trajectory, and then combining pre-implantation with soma. The greatest correlation between developmental stage and pseudotime was found for the second run i.e., running on the two branches separately.
- SCUBA and SLICER algorithms were run using default settings.
- TSCAN (Ji and Ji 2016). To generate pseudotimes we ran TSCAN using the online app. PCA was used for dimensionality reduction, and clusters were selected using the optimal cluster number option. Finally, we manually tuned the ordering of clusters and selected the combination that gave the greatest correlation between developmental stage and pseudotime.
- Wishbone was run using the accompanying GUI, using default settings. Due to numerical issues, we ran with a reduced number of genes compared to that used for Monocle2, TSCAN, SCUBA and SLICER.

Scripts for non-web-based pseudotime methods are available from the GitHub repository: https://github.com/cap76/PGCPseudotime

We evaluated the accuracy of the various approaches by comparing the correlation between the inferred pseudotime order and developmental stage for soma and PGCs separately, and by evaluating a branch-discrepancy metric. Results summarised in Supplementary Figure 1 and Supplementary Table 2. All methods appeared able to broadly separate out the different cells types e.g., pre-implantation, PGCs and soma, but did not necessarily place cells along continuous or consistent trajectories. GPLVM-based approaches offered the best overall performance in terms of branch alignment, followed by Monocle2 and TSCAN.

We also performed GPLVM using an explicit three-branch process. To test that the algorithm was ordering cells in a meaningful way, we checked whether the posterior distribution of pseudotime had diverged from the prior distribution using a Chi-squared test under the null hypothesis that pseudotime was normally distributed with variance defined by the prior. Here we rejected the null hypothesis with a highest p value of <1×10^-23^, indicating that the posterior distribution had diverged from the prior distribution.

## 3 COMPLEX BRANCHING DURING EARLY EMBRYO DEVELOPMENT IN HUMANS

Once a point estimate for the pseudotemporal order of cells had been established, we looked for more complex branch-recombinant structures on a genome scale, by fixing pseudo-times according to the order at step 30,000 in our earlier pseudotime analysis. As the pseudotimes are fixed, this is simply a case of fitting GP model with different branch structures (B-RGP regression). Here we considered the dynamics of PGCs, somatic cells, and ESCs. Each of the three groups was randomly assigned pre-implantation cells with equal probability, except for the ESCs, which partially overlapped with blastocyst stage cells; here we instead randomly assigned pre-blastocyst cells. This assignment reflects our expectation that divergence between PGCs and soma occurs post blastocyst. To capture the dynamics necessary to drive ESCs towards week 4 PGC fate, we also assigned half of the male hPGCs to the ESCs class. For model 1, we assumed:

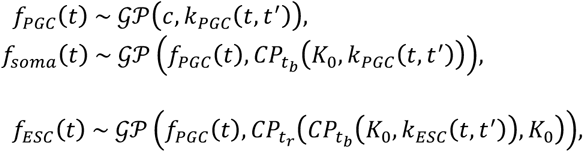

representing genes in which somatic cells diverged from PGCs, and where ESCs diverged from the *in vivo* dynamics of PGC specification before recombining at around week 4. For model 2, we assumed:

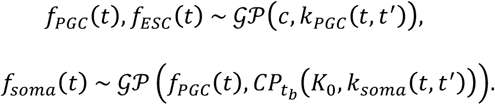

This model represents genes that showed divergence between PGCs and soma, with ESCs identically distributed to PGCs. Finally, for model 3 we assumed:

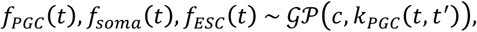

representing genes that showed no divergence between PGCs, soma or ESCs. Because these datasets represent single-cell measurements that might be intrinsically less smooth than the microarray datasets, we assumed a Matérn covariance function throughout. As in previous analyses, we optimised hyperparameters to their MAP values and used the BIC to select the branching structure for each gene. The frequency of the three groups is indicated in Supplementary Figure 9(a), which shows that most genes were not differentially expressed i.e., non-branching. A histogram of the timing of branching is indicated in Supplementary Figure 9(b, c), and shows that of the genes that diverge between PGCs and soma, most do so between blastocyst stage and week 4, as expected.

Following GO analysis on the individual groups using permissive p-values (p<0.1 following multiple hypothesis test correction), we ordered terms by the pseudo-time at which 50% of their associated genes had branched, as previously done in Yang et al. (2016). In Figure 16 we indicate a heatmap representation of this ordering, where the x-axis represents the pseudo-time, the y-axis represents individual GO terms, and the colour indicates the fraction of genes that had branched. From these heatmaps we can see a continuum of responses. An associated table containing the full list of terms associated with the y-axis is available in Supplementary Table 4.

## 4.1. ANALYSIS OF *SOX17* OVEREXPRESSION LINES

We identified differentially expressed genes in hESC lines constitutively over-expressing *SOX17* (Seguin et al. (2008); GSE10809). Differential expression was evaluated using a Student’s t-test on log2 fluorescence versus parental lines using a cut-off of p<0.03 and log2 fold change >2, similar to Seguin et al. (2008).

## 4.2. GENE SET ENRICHMENT ANALYSIS

Gene set based enrichment analysis was used to identify significantly enriched terms using Enrichr database. Here we focused on enrichment of GO 2017 and KEGG 2016 terms using permissive p-values (p<0.1 following multiple hypothesis testing correction). Using Enrichr we also looked for enrichment of Kinase Perturbations from GEO, Single Gene Perturbations from GEO, and Ligand Perturbations from GEO, using p-value cut-off of p<0.05 (following multiple hypothesis testing correction).

## 5 EXPERIMENTAL PROCEDURES

### 5.1 CELL CULTURE AND PGCLC DIFFERENTIATION

hESCs (W15-NANOS3-tdTomato or WIS2-NANOS3-T2A-tdTomato (N3-tdTom, Kobayashi et al. (2017)) were cultured as in (Irie et al. 2015) on irradiated mouse embryonic fibroblasts (MEFs) (GlobalStem) in 4i medium (Supplementary Table 5) or modifications therefrom (Supplementary Table 6). Media were replaced every day. hESCs were passaged by single-cell dissociation using 0.25% Trypsin-EDTA (GIBCO). 10 μM ROCK inhibitor (Y-27632, TOCRIS) was added for 24 hours after passaging.

To induce PGCLCs (Irie et al. 2015), hESCs were trypsinized, filtered and plated into ultra-low cell attachment U-bottom 96-well plates (Corning, 7007) at 4,000 cells/well density in 100 μl PGCLC medium (Supplementary Table 7) The plate was centrifuged at 300g for 3 minutes and placed into a 37°C 5% CO_2_ incubator until embryoid body (EB) collection for downstream analysis. Reporter fluorescence intensities were monitored daily throughout differentiation using Olympus IX71 microscope.

**Supplementary files 3-7 available as separate spreadsheets.**

### 5.2. Flow Cytometry

D4 or D5 EBs were washed in PBS and dissociated with 0.25% Trypsin-EDTA for 15 min at 37°C. Cells were resuspended in FACS buffer (3% FBS in PBS) and incubated with antibodies specified in Supplementary Table 8. After washing with FACS buffer, the cells were recorded on BD LSR Fortessa. Data were analysed using FlowJo (Tree Star).

### 5.3. Real-Time Quantitative RT-PCR

Total RNA was extracted from unsorted hESCs using RNeasy Mini Kit (QIAGEN). cDNA was synthesized using QuantiTect Reverse Transcription Kit (QIAGEN). qPCR was performed on a QuantStudio 12K Flex Real-Time PCR machine (Applied Biosystems) using SYBR Green JumpStart Taq ReadyMix (Sigma) and human-specific primers (Supplementary Table 9). The ∆∆*C*t method was used for quantification of gene expression. Three technical replicates were used for each biological replicate.

### 5.4. Dominant-negative ERK expression

For inducible expression of dominant-negative ERK1 (DN-ERK), a cDNA from pFLAG-CMV-hErk1(K71R) (Addgene plasmid #49329) encoding human ERK1 with K71R mutation (Robbins et al. 1993) was cloned (into a doxycycline (dox)-inducible PB-TRE-3G vector (Kobayashi et al. 2017) to yield pPB-TRE-3G-DN-ERK-IRES-EGFP. The plasmid was generated using In-Fusion cloning (Clontech) according to manufacturer’s recommendations. Primers used for cloning are specified in Table 6f. N3tdTom hESCs were co-lipofected with 2 μg PB-TRE-3G-DN-ERK-IRES-EGFP, 2.5 μg PBase and 0.5 μg pPBTET3G-Neo (Kobayashi et al. 2017). Lipofection was performed using OptiMEM (GIBCO) and Lipofectamin 2000 (Invitrogen) according to manufacturer’s recommendations.

---

**Supplementary Figure 1:**
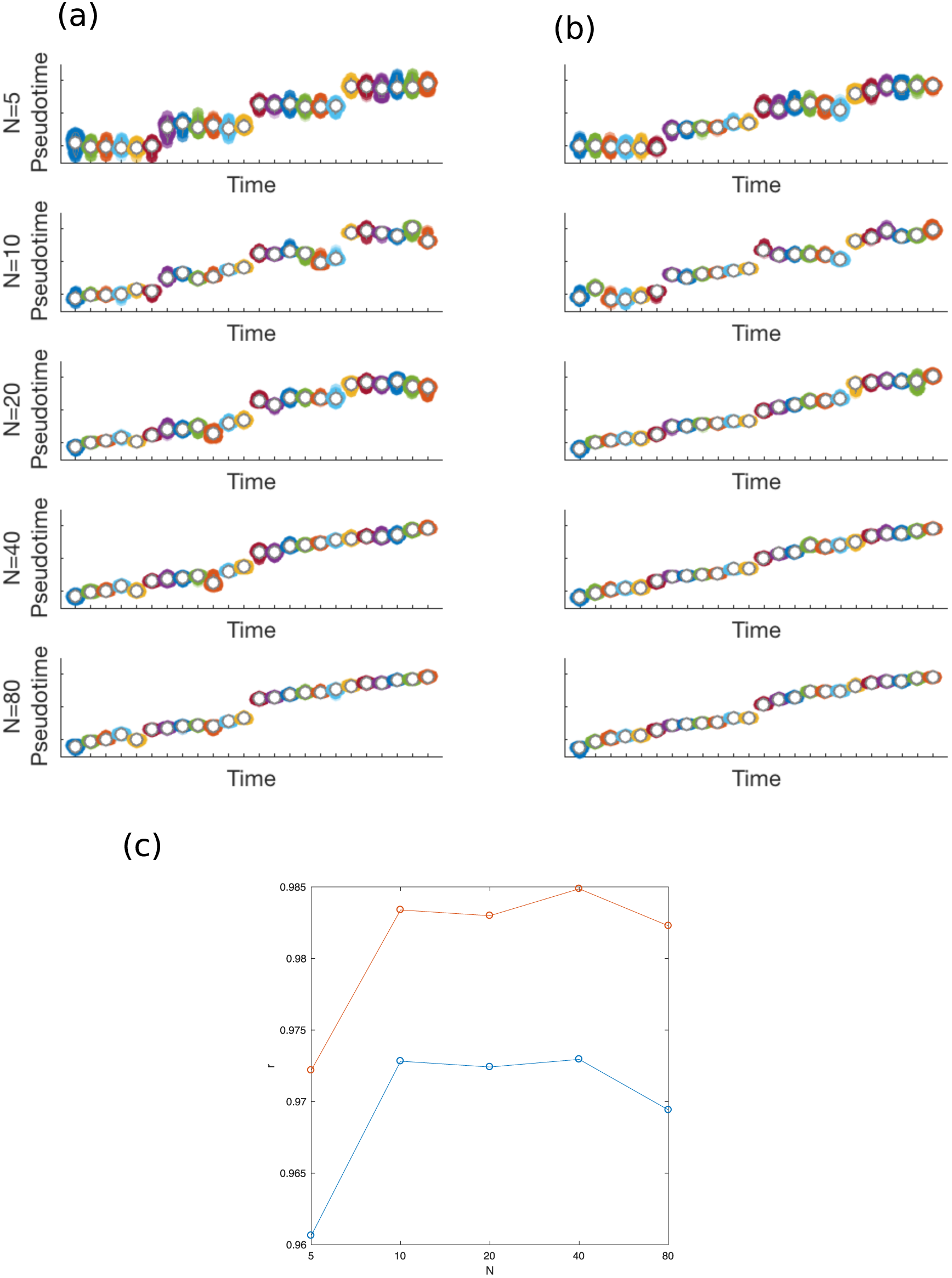
The pseudotemporal order (y-axis) is plotted against the true order (x-axis) for *Arabidopsis thaliana* transcriptional data for the control time-series (a) and *Botrytis cinerea* infected time series (b). (c) The mean correlation (over 5 random initialisations) between time and pseudotime as the number of marker genes increases.

**Supplementary Figure 2:**
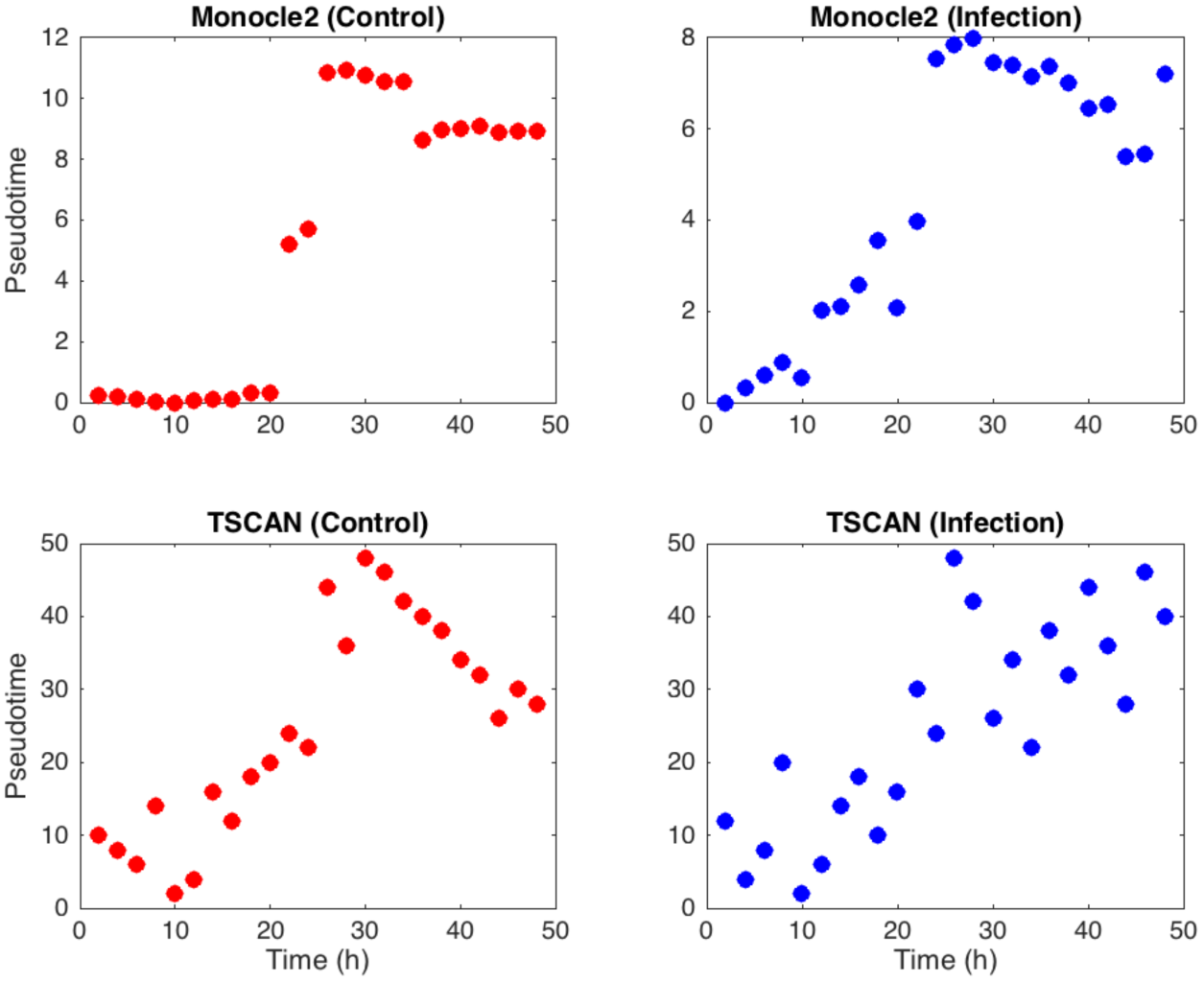
The pseudotemporal order (y-axis) is plotted against the true time (x-axis) for *Arabidopsis thaliana* transcriptional data using Monocle2 (Top) and TSCAN (Bottom) respectively.

**Supplementary Figure 3:**
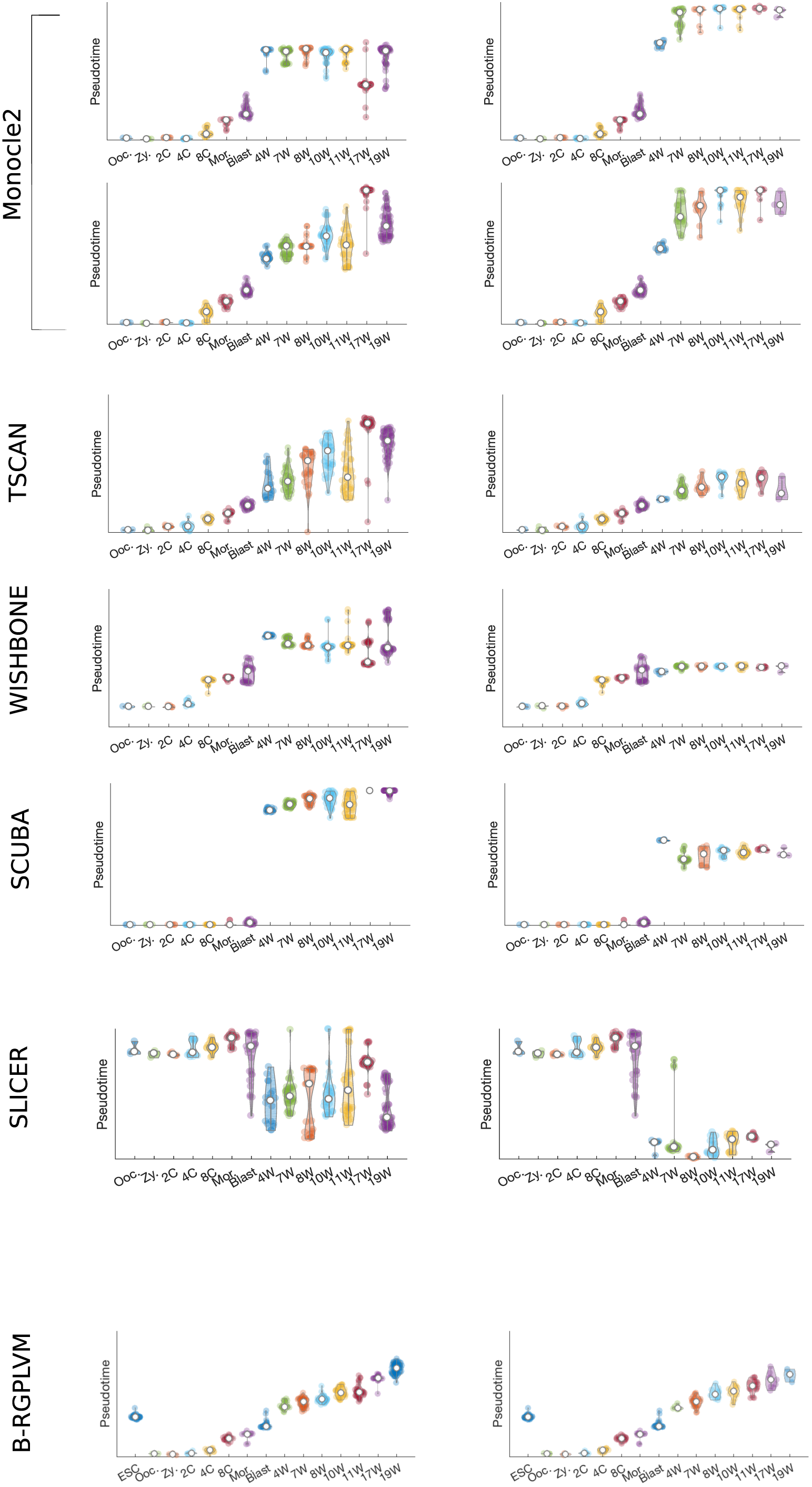
Comparison of pseudotime ordering using MONOCLE, TSCAN, Wishbone, SCUBA and SLICER versus B-RGPLVM. Here we indicate the inferred pseudotime (y-axis) versus the true developmental stage. All approaches appear to be able to generally separate out pre-implantation cells from either PGCs or soma, with B-RGPLVMs performing best. Together, these observations highlight the increased accuracy afforded when including prior information about capture time in B-RGPLVMs and a more realistic generative model in terms of an underlying branching process. Note that the distribution of cells from pre-implantation embryos will be identical when comparing within a method.

**Supplementary Figure 4:**
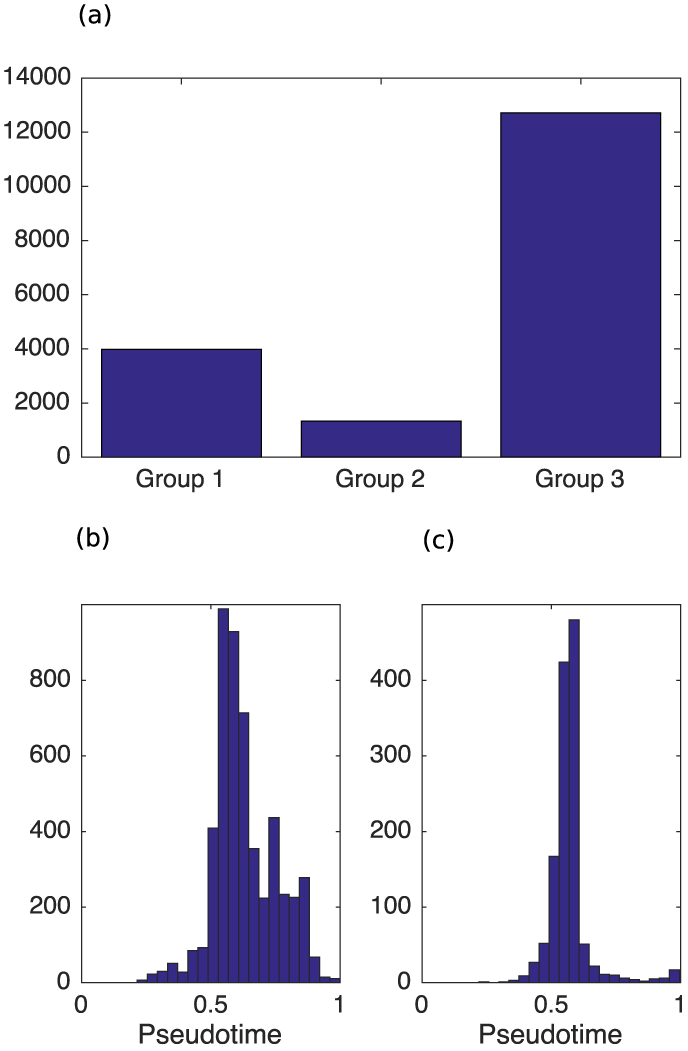
**(a)** Frequency of the different groups. **(b)** Timing of branching between PGC and somatic cells. **(c)** Time of recombination between ESCs and inferred *in vitro* dynamics.

**Supplementary Figure 5:**
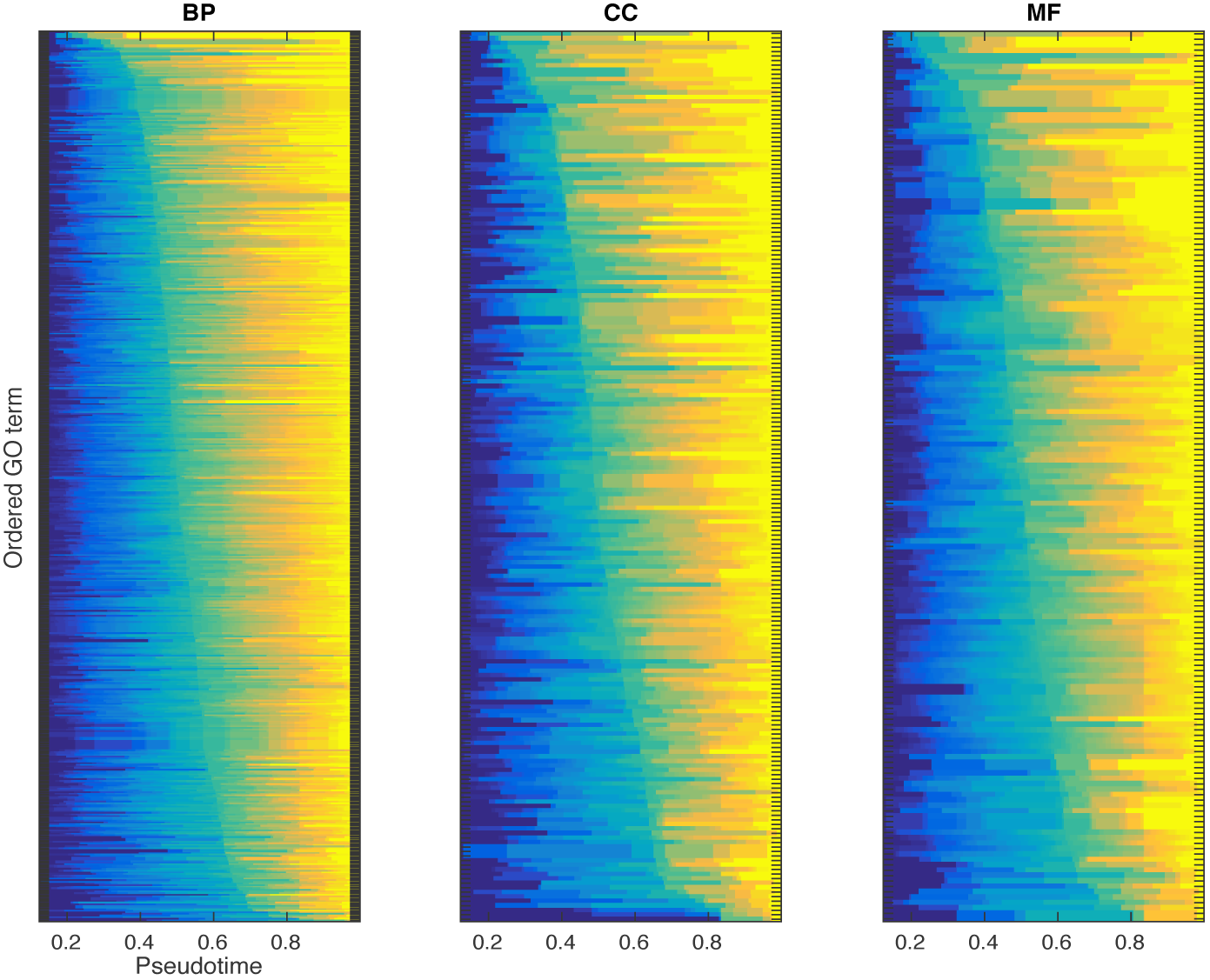
Individual GO terms for genes that were up-regulated in PGCs versus soma, according to the timing at which at least 50% of the genes associated with each term had branched. See also associated Supplementary Table 4.

**Supplementary Figure 6:**
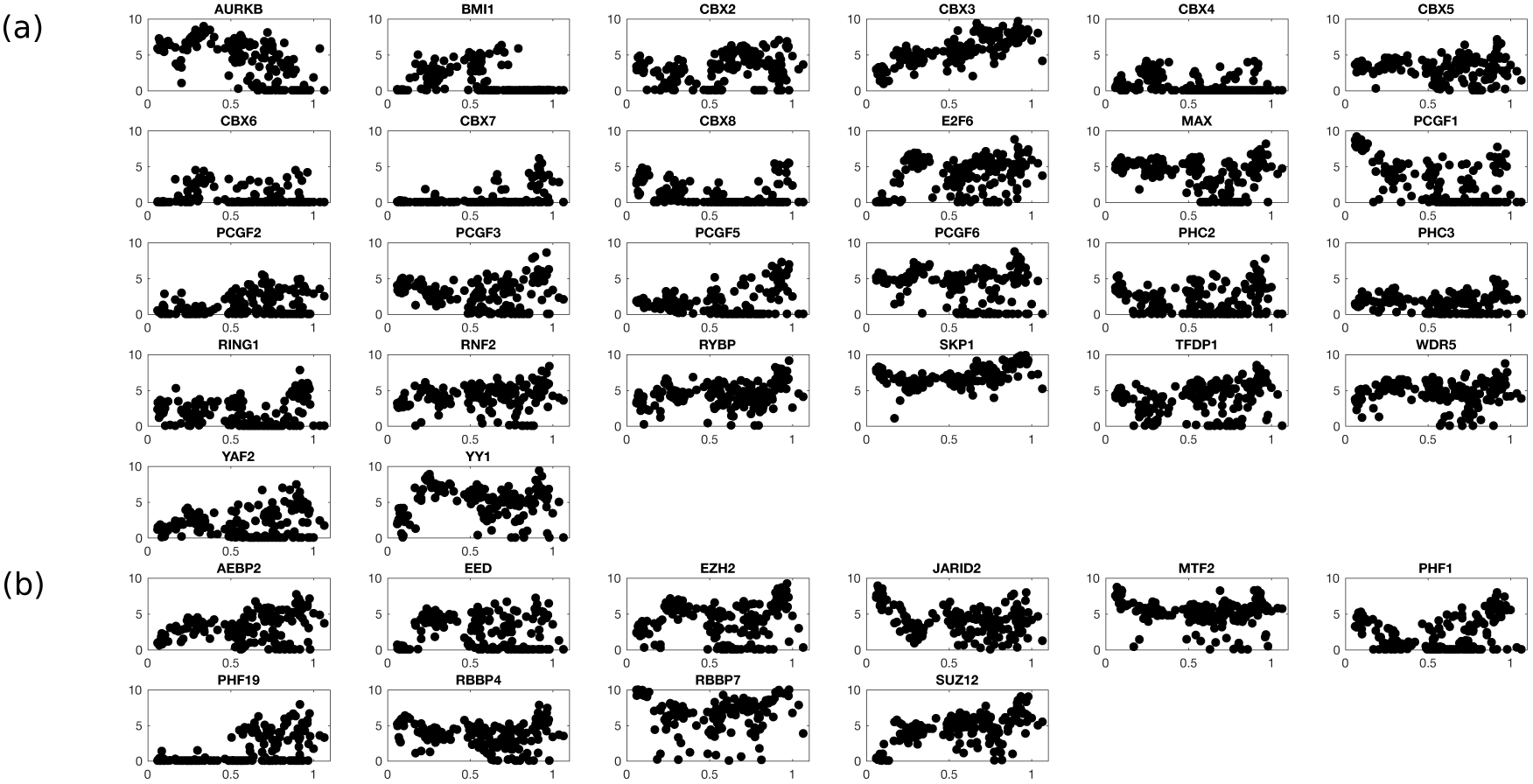
Pseudotime trajectories for male PGCs for (a) polycomb repressive complex 1 (PRC1) and (b) polycomb repressive complex 2 (PRC2) reveal highly dynamic changes during development.

**Supplementary Figure 7:**
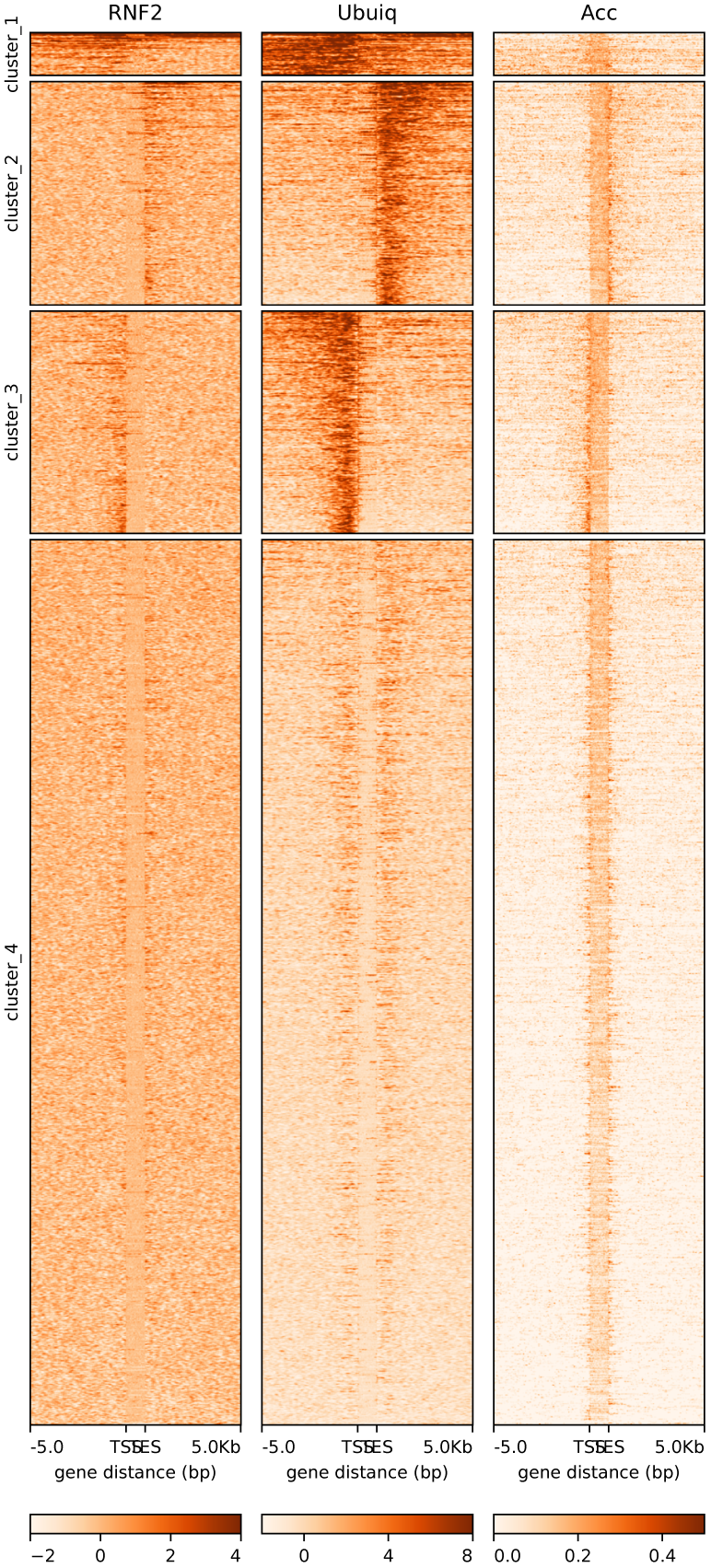
The clustering of early-branched genes by the level of RNF2 and H2AK119ub levels in H1 ESCs (over gene bodies +/- 5kb). Crucially these sites remain accessible during PGC development, as indicated by chromatin accessibility (NOME-seq) in week 11 male hPGCs (Guo et al. 2017).

**Supplementary Figure 8:**
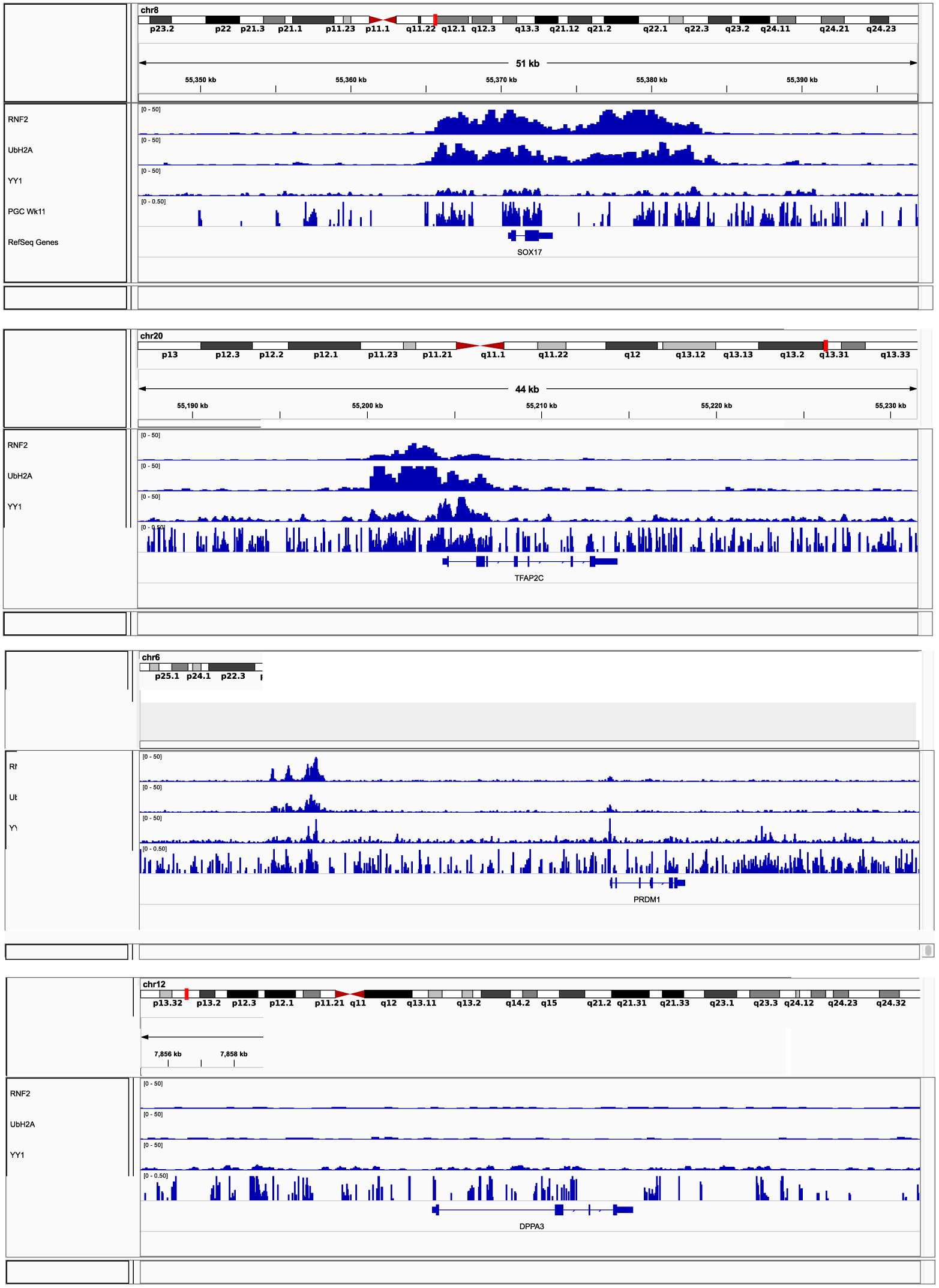
Gene browser track for RNF2, H2AK119ub and YY1 at several PGC-related genes, SOX17, TFAP2C, PRDM1, and DPPA3. The H2AK119ub mark was found at SOX17 and TFAP2C in H1 ESCs, but not at PRDM1 or STELLA. YY1 has been shown to bind SOX17, TFAP2C and PRDM1, but not STELLA, in HEK293 cells. Crucially, several key putative targets of RNF2 and YY1 were shown to be accessible in week 11 male hPGCs (Guo et al. 2017)

**Supplementary Figure 9:**
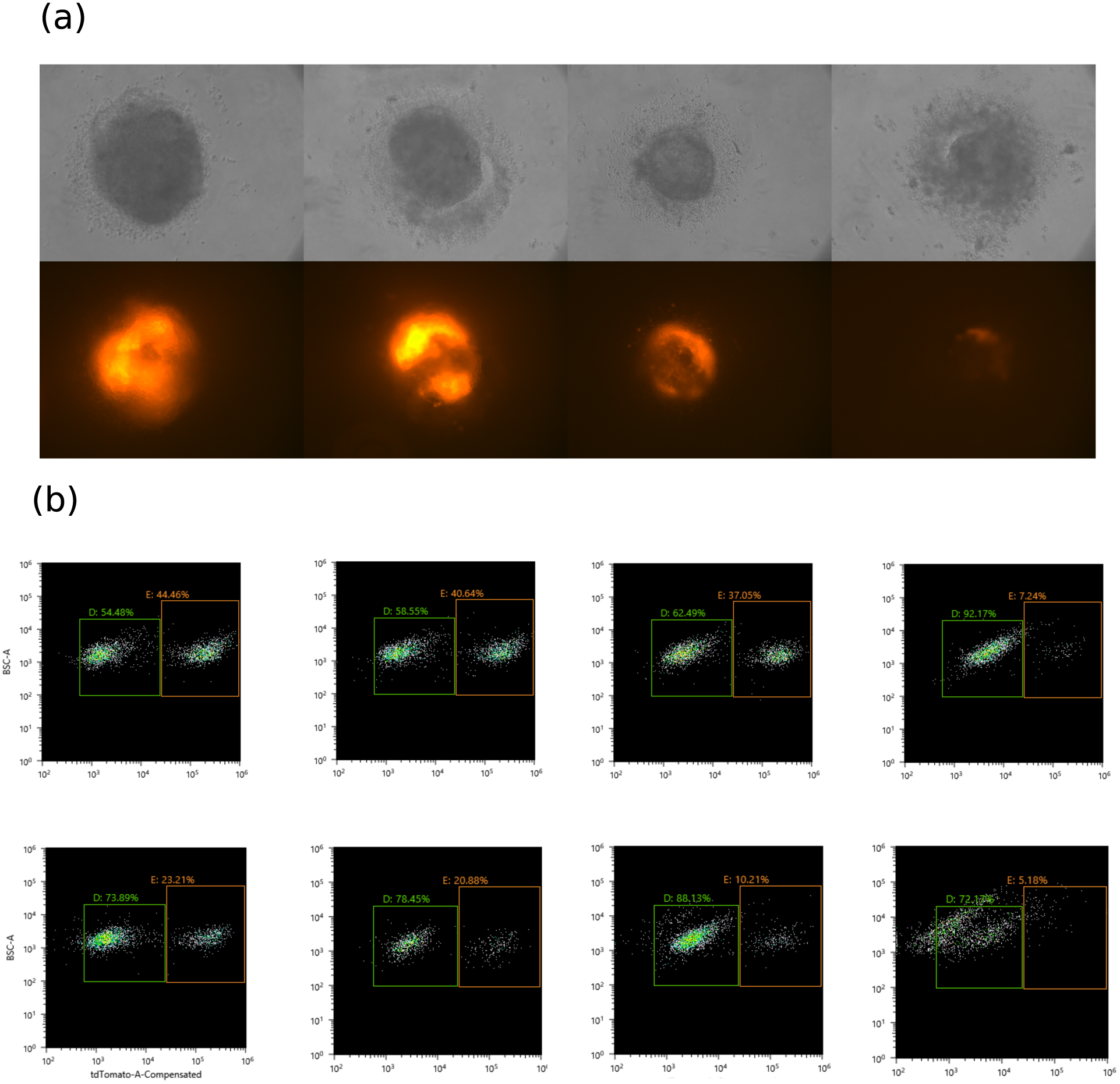
W15-NANOS3-tdTomato reporter line in in PGC media and following increase dose PRC1 RING1B/BMI1-dependent ubiquitination inhibitor PRT 4165 (12μM, 25μM, 50μM). (b) FACS quantification in two independent replicates: (middle row) control, 12μM, 25μM, 50μm; (bottom row) control 30μM, 40μM, 50μM.

**Supplementary Figure 10:**
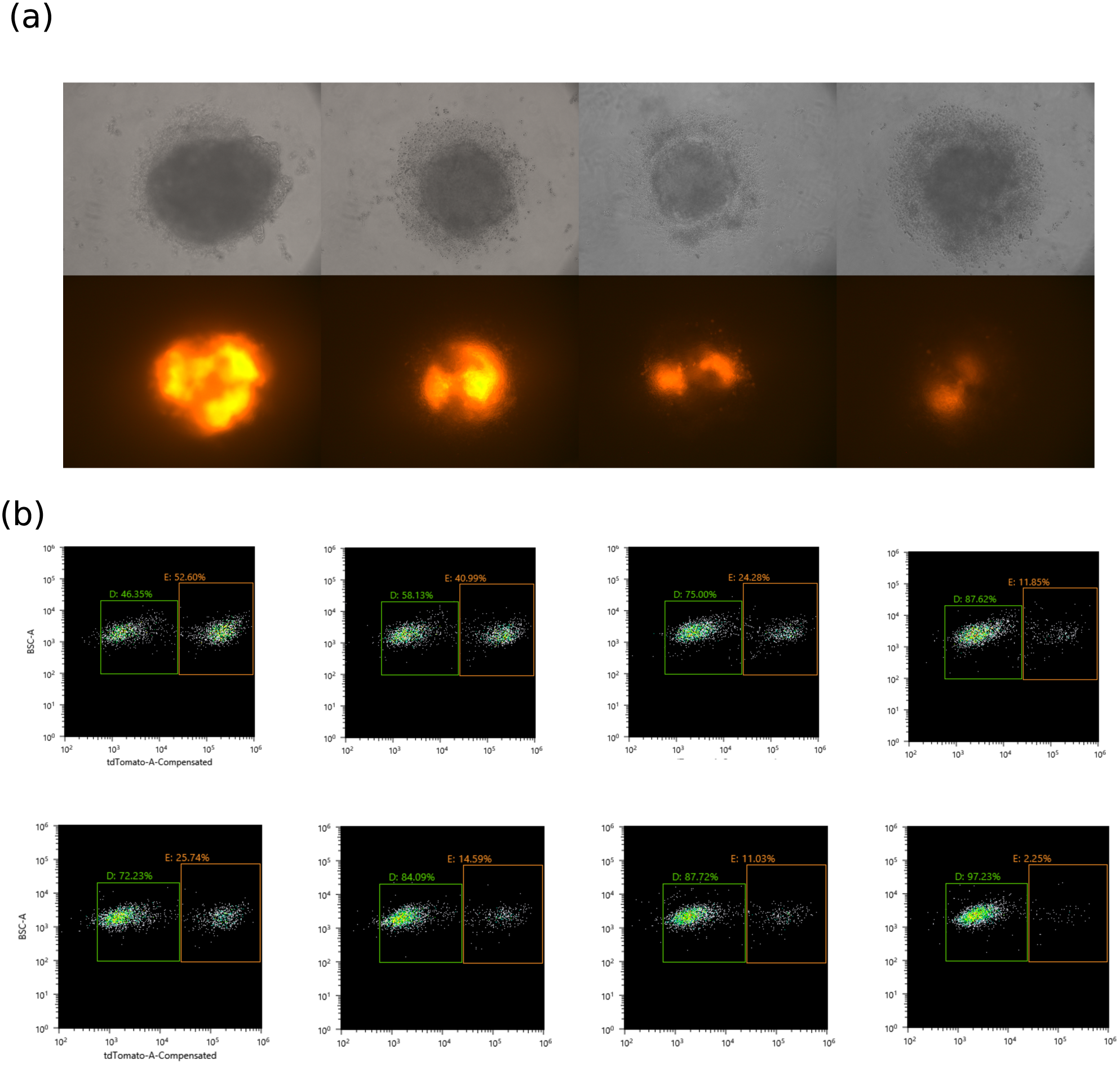
W15-NANOS3-T2A-tdTomato reporter line in PGC media and following increase dose with small molecule inhibitor DETA/NONOate. (b) FACS quantification in two independent replicates: (middle row) control, 0.1mM, 0.5mM and 1mM; (bottom row) 0.25mM, 0.5mM, 1mM.

**Supplementary Figure 11:**
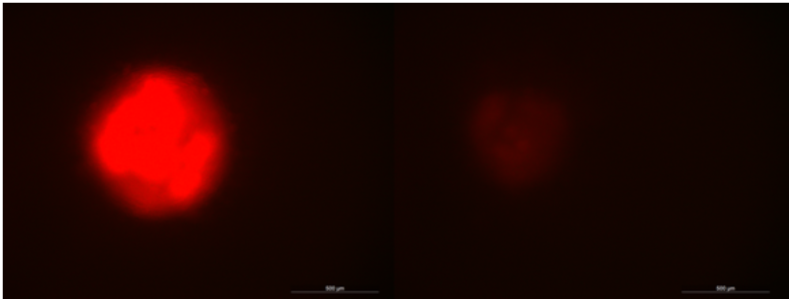
W15-NANOS3-tdTomato in PGC media (left) and with EZH2-mediated histone methylation inhibitor DZNep (1μM).

**Supplementary Figure 12:**
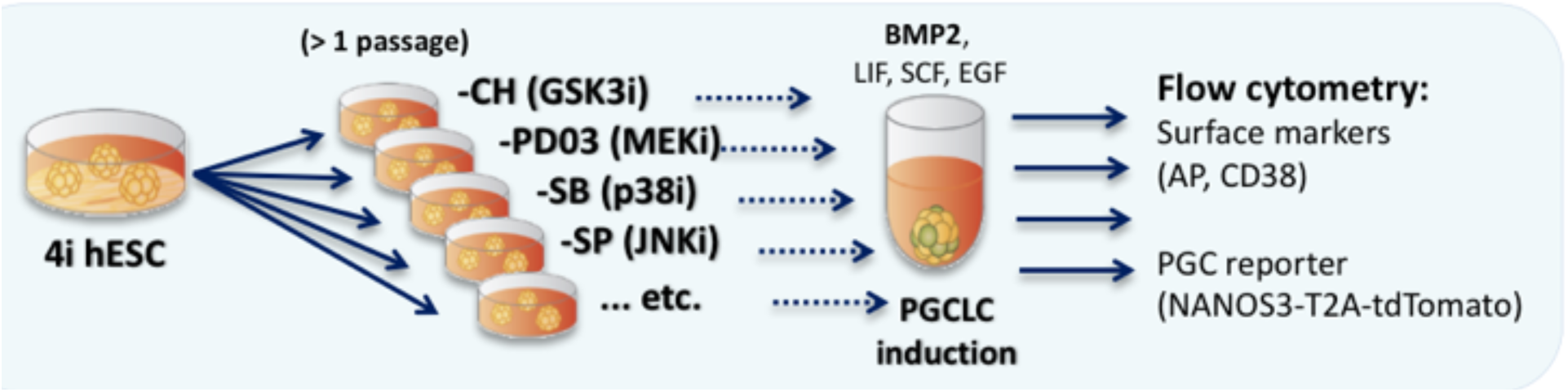
Scheme of the experimental workflow used to test the roles of individual components of the 4i hESC medium to sustain PGCLC competence. hESCs cultured in 4i medium were transferred to media lacking one of the kinase inhibitors for at least one passage. These cells were then subjected to standard PGCLC induction with BMP2 and supporting cytokines. PGCLC induction efficiency was assessed by flow cytometry as percentage of live cells expressing PGC markers AP, CD38 and NANOS3. Changes in PGCLC induction efficiency were thus used as proxy of changes in competence for PGC fate.

**Supplementary Figure 13.**
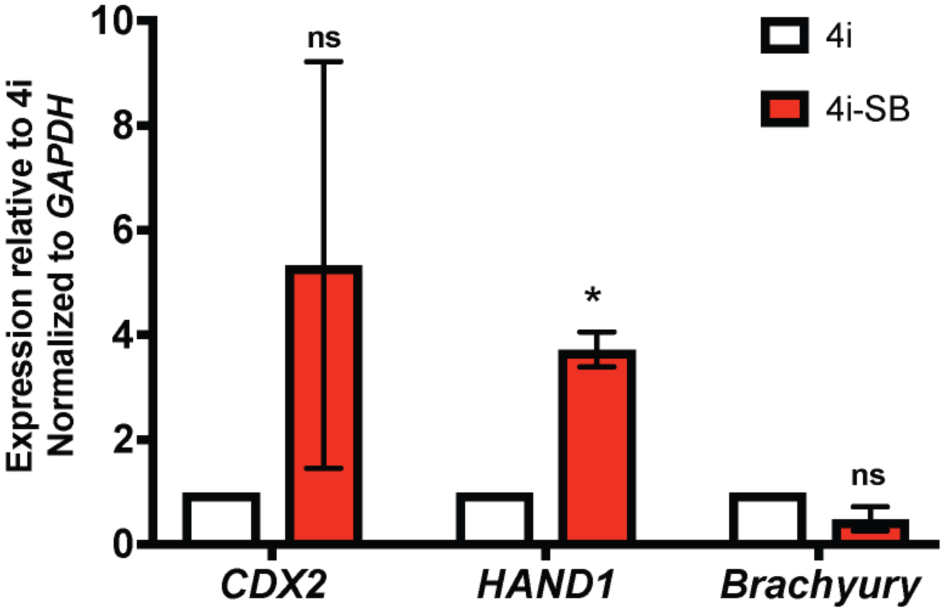
Withdrawal of p38 inhibitor from 4i results in hESC differentiation. hESCs were cultured in respective media for 4 passages and collected for qPCR analysis. Data are shown as mean (of three technical replicates each) ± SD of 2 independent experiments. * p ≤ 0.05, ns: not significant (p > 0.05), Holm-Sidak t-test.

**Supplementary Figure 14.**
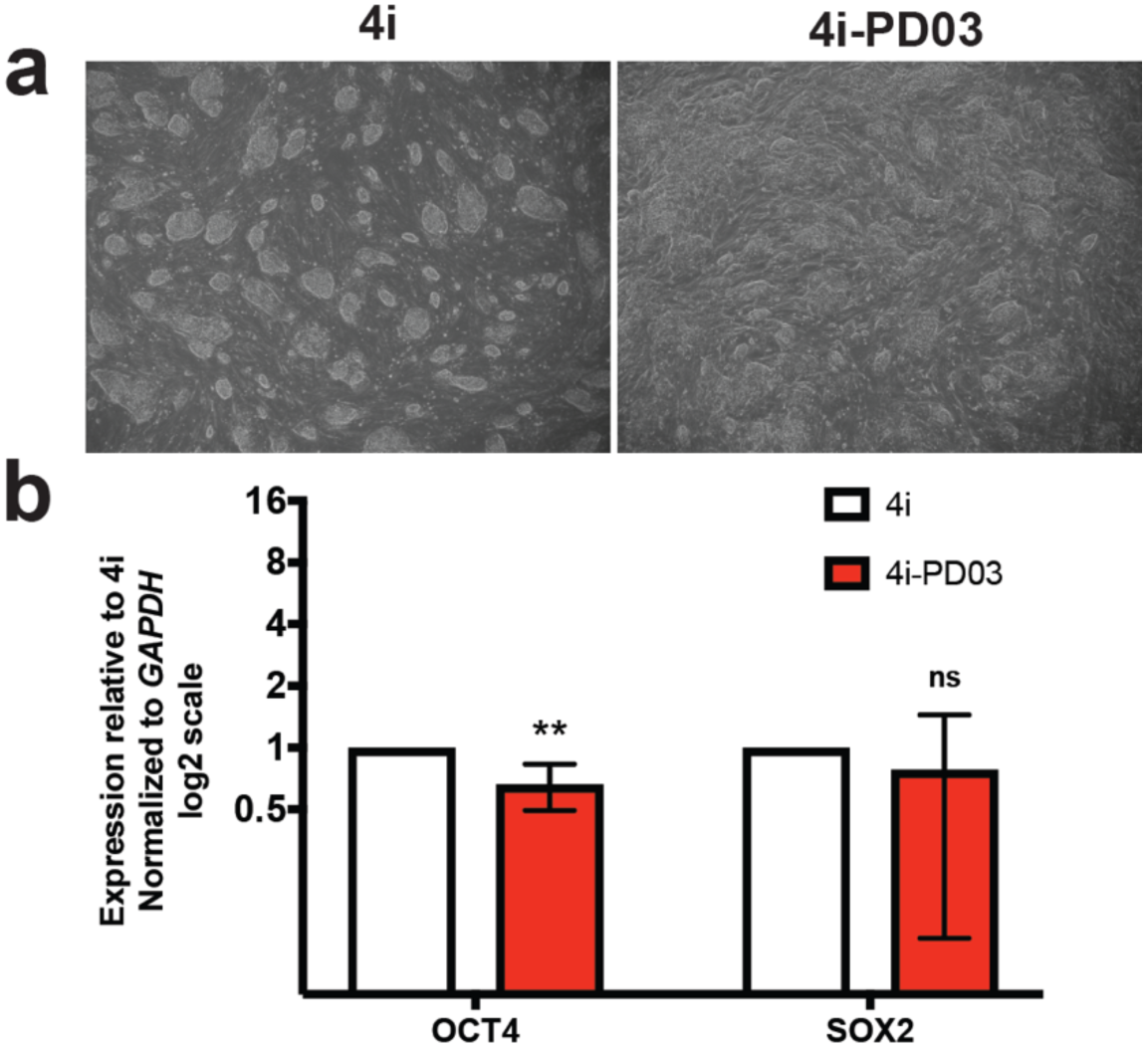
Withdrawal of MEK inhibitor from 4i results in changes to hESC colony morphology and induces expression of endoderm-related markers. (**a)** Representative pictures of hESCs cultured in complete 4i or “4i-PD03” media. **(b)** qPCR showing pluripotency and endoderm marker genes expression in bulk hESCs cultured in 4i and “4i-PD03” media. Data are shown as mean (of three technical replicates each) ± SD of 5 independent experiments. * p ≤ 0.05, ** p ≤ 0.01, ns: not significant (p > 0.05), Holm-Sidak t-test.

**Supplementary Figure 15.**
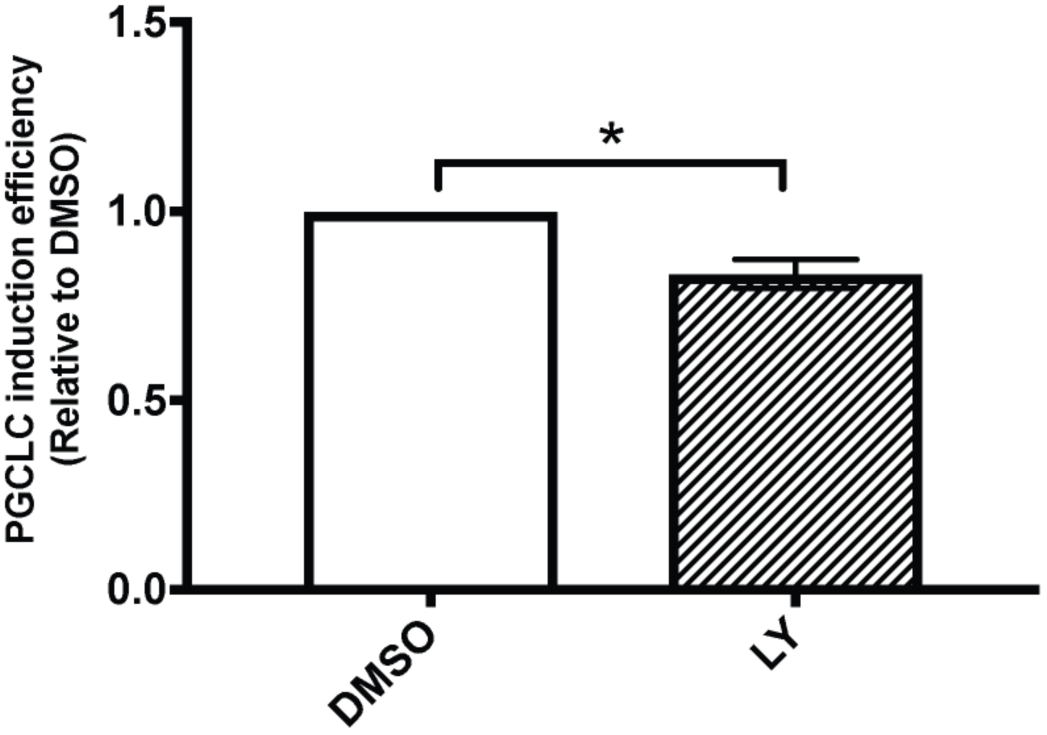
Inhibition of PI3K during differentiation does not abrogate PGCLC specification. Data are shown as mean ± SD of 2 independent experiments. * p ≤ 0.05, Holm-Sidak t-test.

**Supplementary Table 1.** Number of cells for each cell type

| Stage | $N$ | Source |
| --- | --- | --- |
| ESCs | 34 | Yan et al. (2013) |
| Pre-implantation | 90 | Yan et al. (2013) |
| PGCs | 242 | Guo et al. (2015) |
| Soma | 86 | Guo et al. (2015) |

**Supplementary Table 2.** Benchmarking of pseudotime algorithms on the PGC data

| Stage | R (PGC) | R (soma) | $\Delta_b$ |
| --- | --- | --- | --- |
| GPLVM (2-branch) | 0.96 | 0.95 | 0.004 |
| GPLVM (3-branch) | 0.98 | 0.97 | **0.002** |
| GPLVM (3-branch) | 0.97 | 0.96 | 0.006 |
| GPLVM (3-branch) | **0.99** | **0.99** | 0.003 |
| GrandPrix | 0.84 | 0.87 | 0.02 |
| MONOCLE2 | 0.84 | 0.87 | 0.35 |
| SCUBA | 0.85 | 0.85 | 1.1 |
| SLICER | -0.50 | -0.80 | 1.1 |
| TSCAN | 0.85 | 0.92 | 0.44 |
| Wishbone | 0.74 | 0.80 | 0.48 |

**Supplementary Table 5.**
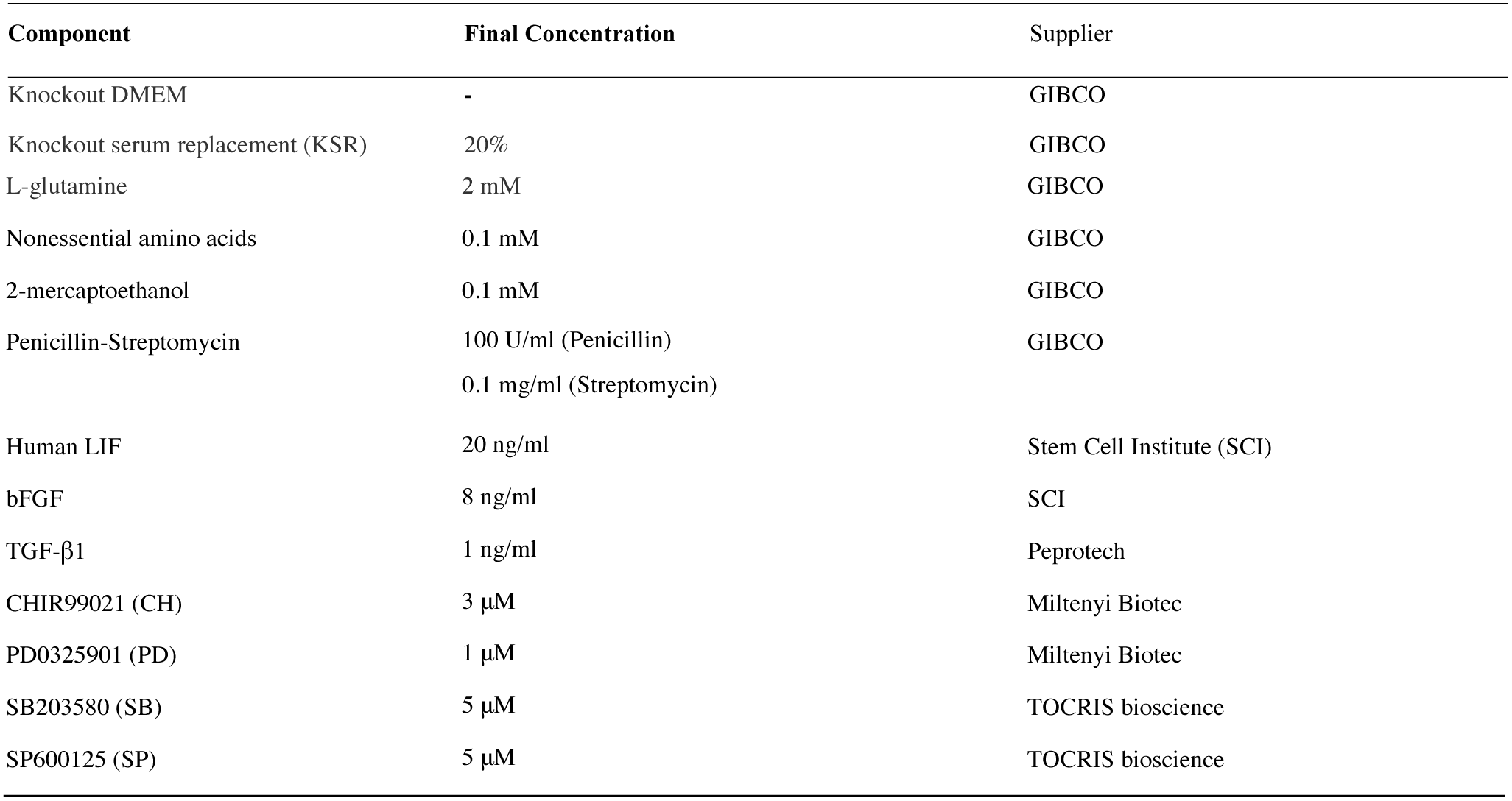
Competent (4i) medium composition.

**Supplementary Table 6.** Modifications of the 4i medium used in the competence screen.

| Component | Final Concentration |
| --- | --- |
| 4i-PD03 | 4i as in Supp. Table 5 without PD0325901 |
| 4i+PD17 | 4i as in Supp. Table 5 supplemented with indicated concentrations of PD173074 (TOCRIS bioscience) |
| 4i-PD03+PD17 | 4i-PD as above supplemented with indicated concentrations of PD173074 (TOCRIS bioscience) |
| 4i+LY | 4i as in Supp. Table 5 supplemented with 10 μM LY294002 (Sigma) |
| 4i-PD+LY | 4i-PD as above supplemented with 10 μM LY294002 (Sigma) |
| 4i+IWR | 4i as in Supp. Table 5 supplemented with 2.5 μM IWR-1 (Sigma) |
| 4i-CH | 4i as in Supp. Table 5 without CHIR99021 |
| 4i-CH+IWR | 4i as in Supp. Table 5 without CHIR99021 supplemented with 2.5 μM IWR-1 (Sigma) |
| 4i-SB | 4i as in Supp. Table 5 without SB203580 |
| 4i-SP | 4i as in Supp. Table 5 without SP600125 |
| 4i-LIF | 4i as in Supp. Table 5 without human LIF |
| 4i+JAKi | 4i as in Supp. Table 5 supplemented with 1μM CAS457081-03-7 |

**Supplementary Table 7.** PGCLC induction medium composition

| Component | Final Concentration | Supplier |
| --- | --- | --- |
| Glasgow's MEM (GMEM) | - | GIBCO |
| KSR | 15% | GIBCO |
| L-glutamine | 2 mM | GIBCO |
| Nonessential amino acids | 0.1 mM | GIBCO |
| 2-mercaptoethanol | 0.1 mM | GIBCO |
| Penicillin-Streptomycin | 100 U/ml (Penicillin)<br>0.1 mg/ml (Streptomycin) | GIBCO |
| Sodium pyruvate | 1 mM | Sigma |
| BMP2 | 500 ng/ml | SCI |
| Human LIF | 1 µg/ml | SCI |
| SCF | 100 ng/ml | R&D Systems |
| EGF | 50 ng/ml | R&D Systems |
| ROCK inhibitor (Y-27632) | 10 µM | TOCRIS bioscience |

**Supplementary Table 8.**
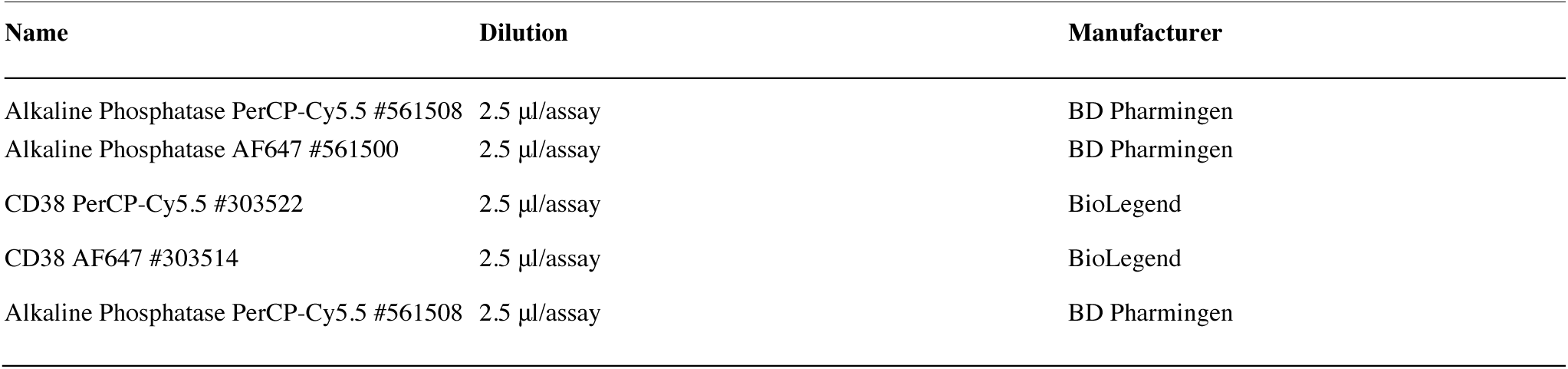
Antibodies used for flow cytometry

| Name | Dilution | Manufacturer |
| --- | --- | --- |
| Alkaline Phosphatase PerCP-Cy5.5 #561508 | 2.5 µl/assay | BD Pharmingen |
| Alkaline Phosphatase AF647 #561500 | 2.5 µl/assay | BD Pharmingen |
| CD38 PerCP-Cy5.5 #303522 | 2.5 µl/assay | BioLegend |
| CD38 AF647 #303514 | 2.5 µl/assay | BioLegend |
| Alkaline Phosphatase PerCP-Cy5.5 #561508 | 2.5 µl/assay | BD Pharmingen |

**Supplementary Table 9.**
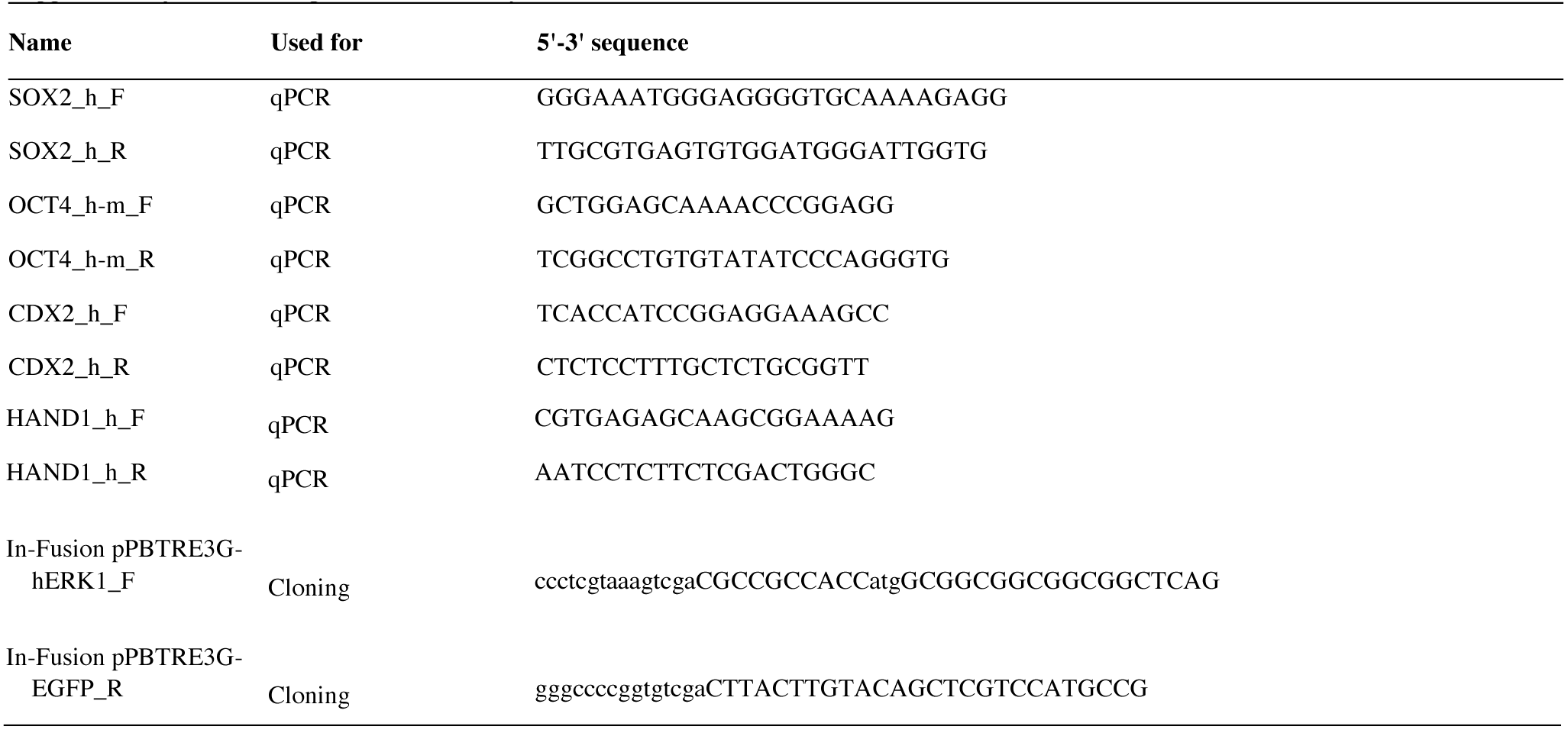
Oligos used in the study.

